# Ecological inference from isolated vertebrae: Evaluating functional signal across the carnivoran spine

**DOI:** 10.1101/2025.08.16.670649

**Authors:** Julia A. Schwab, Borja Figueirido, Katrina E. Jones

**Affiliations:** Oxford University Museum of Natural History, University of Oxford, Parks Rd, OX1 3PW Oxford, United Kingdom; Department of Earth and Environmental Sciences, University of Manchester, M13 9PL Manchester, United Kingdom; Departamento de Ecología y Geología, Facultad de Ciencias, Campus de Teatinos s/n, Universidad de Málaga, 29071-Málaga, Spain; School of Earth Sciences, University of Bristol, Bristol BS8 1RJ, UK

**Keywords:** Ecomorphology, Locomotion, Carnivora, Vertebral coloumn, Geometric morphometrics

## Abstract

Understanding the ecological adaptations of extinct species is a central goal in vertebrate palaeontology, but is often limited by the incomplete nature of the fossil record. While skulls and limb bones have traditionally been emphasized in functional and ecological reconstructions, vertebrae are frequently overlooked. While isolated vertebrae are among the most commonly preserved postcranial elements, they are rarely found as complete vertebral columns, raising the question of whether isolated elements alone can yield meaningful ecological information. In this study, we assess the potential of vertebral morphology to predict two key ecological traits, running speed and hunting mode, using three-dimensional geometric morphometrics across ten presacral vertebrae from a broad sample of extant carnivorans. We evaluate the predictive power of individual vertebrae, regional groupings (cervical, thoracic, lumbar), and multi-element combinations. Our results show that certain vertebrae retain strong ecological signals on their own, especially the first thoracic and lumbar elements. However, combining multiple vertebrae often dilutes ecological signal, likely due to their differing functional roles along the axial column. This highlights the importance of treating vertebral regions independently and suggests that single, strategically informative vertebrae may outperform multi-element approaches in some contexts. We apply this framework to the extinct dire wolf (*Canis dirus*) and find contrasting signals along the spine, the first thoracic and lumbar vertebrae suggest adaptations for faster locomotion, while some cervical vertebrae indicate an intermediate running speed. This mosaic supports the idea that *C. dirus* occupied a complex ecological niche involving both active predation and scavenging. These findings underscore the power of vertebral morphology for ecological inference in fossil taxa, particularly when remains are fragmentary, and argue for a more nuanced use of isolated axial elements in reconstructing extinct carnivoran behaviour.

## 1 Introduction

Inferring ecology from the skeletal morphology of extinct species is a major challenge for paleobiologists trying to understand the evolution of ancient ecosystems. When studying the fossil record, most studies focus on either isolated elements (e.g. skull, ankle), which are homologous across species or they focus on multiple elements from a single, well-preserved, specimen. However, studying serial elements, such as the vertebrae, in fossil taxa presents an even greater challenge. Firstly, homologizing among elements and across taxa with different vertebral counts is difficult. Secondly, complete vertebral columns with all elements preserved are rare, which leaves only isolated elements to study. This raises the question if single vertebrae can be useful when drawing inferences about the ecology and lifestyle in extinct taxa, and further, which specific elements may be most valuable. Further, where multiple isolated elements are preserved, to what extent might combining information from them improve our ecological inferences over single element analyses, and which combinations may be best? To address these questions, we explored the ecological signal across the vertebral column of carnivorans.

Carnivora are one of the most diverse mammal groups, occupying a wide range of ecological niches and habitats. They originated in the Palaeocene, about 65 million years ago (Janis et al. 1998), and have become specialised for a wide range of locomotor modes, hunting strategies, and ecologies. Pursuit predators are usually large species that chase their prey over long distances and engage in a high-speed terrestrial locomotion, like the wolf (*Canis lupus*) or the cheetah (*Acinonyx jubatus*). Whereas ambush hunters stalk their prey before a short chase like most felids do (Van Valkenburgh 1985). Smaller species often pounce on their prey, but some species rarely hunt at all, for example slower-moving herbivorous or insectivorous taxa such as the red panda (*Ailurus fulgens*) or the badger (*Meles meles*). These specializations have impacted limb (e.g., Andersson 2004; Martín-Serra et al. 2014; Figueirido et al. 2015; Figueirido et al. 2023) and cranial morphology (e.g., Figueirido et al. 2010, 2011, 2013; Grohé et al. 2018; Schwab et al. 2019; Lyras et al. 2023), which can be diagnostic in inferring locomotory or dietary ecology in the fossil record. However, relatively little is known about the locomotor specializations of the vertebral column, nor its power to discern locomotor function in extinct species.

The morphology of the vertebral column correlates with locomotor mode and ecology in mammals more broadly (e.g., Kort and Polly 2023; Belyaev et al. 2023; Kort and Jones 2024), and in specific carnivoran groups (e.g., Jones 2015; Figueirido et al. 2021; Belyaev et al. 2024). Further, the vertebral column can be subdivided into regions, and a better understanding of spinal regionalisation can shed light on the impact of functional modules on locomotor adaptation (e.g., Randau et al. 2018; Martín-Serra et al. 2021; Belyaev et al. 2023; Gillet et al. 2024; Taewcharoen et al. 2024). In terrestrial mammals, the vertebral column can be divided into three different regions based on their morphology and function, the cervical region (neck vertebrae), the thoracic region (associated with rib attachments) and the lumbar region (most dorsal presacral region). The last two regions are mostly associated with locomotion in mammals and have been associated with running patterns (e.g., Galis et al. 2014; Belyaev et al. 2022, 2023). In felids, the vertebral column has been associated with locomotor mode, and morphology of the lumbar region is particularly useful for differentiating between arboreal, scansorial and terrestrial species (Randau et al. 2016; Figueirido et al. 2021). However, different prey killing techniques have not directly been linked to lumbar morphology (Jones 2015) and can be better associated with forelimb and cranial morphology than with the vertebral column (Slater and Van Valkenburgh, 2009; Meachen-Samuels and Van Valkenburgh, 2009a, b).

Here we examine vertebral morphology in a wide range of extant carnivoran taxa (Figure 1), to test their utility in drawing inferences on running speed and hunting behaviour. Using three-dimensional geometric morphometrics (3D GM), we tested for correlations of running speed and locomotor mode with ten vertebrae distributed along the presacral series. We assess the predictive power of vertebral regions (cervical, thoracic, lumbar), individual vertebrae, as well as different combinations of multiple vertebrae, for these ecological variables. Establishing this analytical framework for ecomorphological analysis of vertebrae will improve the understanding of the palaeobiology of fossil species, where only limited vertebral elements tend to be preserved. We predict that lumbar vertebrae will provide the most ecological signal, since this region has been linked with locomotion, and that addition of more vertebrae will generally increase the ecological signal. Finally, we apply this framework to generate ecological predictions for a well-known extinct carnivoran (Canis dirus) based on isolated fossil vertebrae, to test whether vertebral morphology supports the widely held view that C. dirus was a fast pursuit predator adapted to relatively open habitats, while also evaluating competing hypotheses that it may have been a specialized bone-crusher, a pack-hunting cursorial predator, or a flexible generalist with mixed foraging strategies.

**Figure 1.**
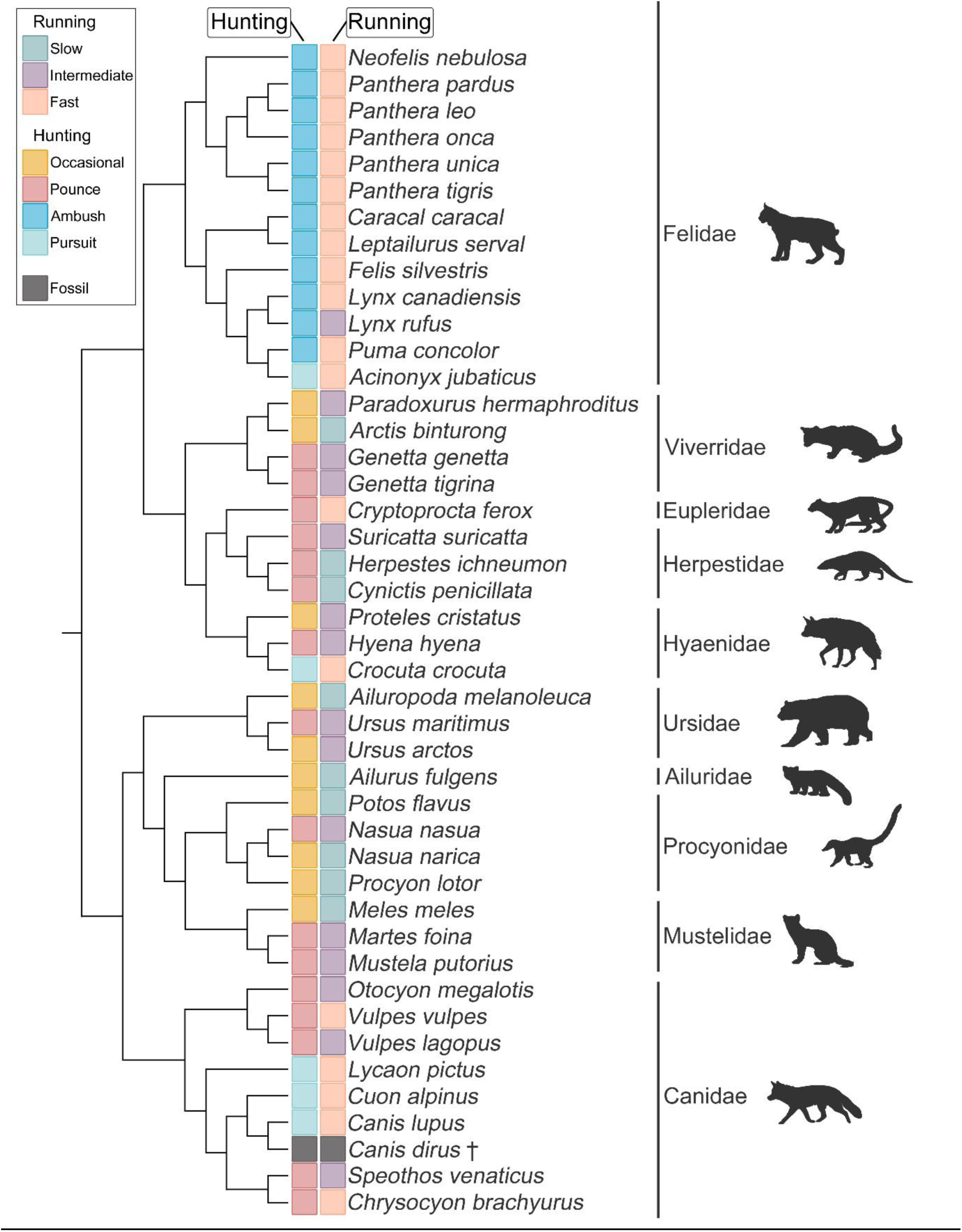
Simplified phylogeny of analysed extant taxa and the extinct *Canis dirus*, including categorisation of their running speed and hunting behaviour (based on Van Valkenburgh et al., (1985 and Galis et al., 2014).

## 2 Material and Methods

### 2.1 Dataset

We sampled vertebral columns from of a total of 43 extant species (52 specimens) of terrestrial carnivorans, belonging to 10 different families (see Suppl. Table 1 for full specimen details). Presacral vertebrae of 13 taxa were scanned with a surface scanner (Artec Spider handheld scanner) to obtain three-dimensional models, which were then used for geometric morphometric analysis. An additional 39 specimen were obtained from Figueirido et al. (2021) and included into the analysis. Additionally, we added vertebrae of the Pleistocene canid *Canis dirus* (see Suppl. Table 2). *C. dirus* been selected as it is one of the most abundant and well-preserved fossil carnivorans, with extensive postcranial material including vertebrae. As a close relative of modern wolves, it provides a valuable case study for testing whether vertebral morphology can reconstruct ecological traits such as running speed and predation strategy in extinct species.

To account for variation in the number of vertebrae for each region (cervical, thoracic, lumbar) in different carnivoran taxa, we have subsampled the vertebral column to ten homologous vertebral positions. We sampled the first, middle and last vertebrae for each region, and for taxa with even numbers we took the vertebra after the middle one. For the cervical region we excluded C1 and C2 due to their unique morphology, and sampled C3, C5 and C7 in the analysis. For the thoracic region, we additionally sampled the diaphragmatic vertebrae. Throughout the text we refer to different vertebrae and combinations as follows: first cervical, CF; middle cervical, CM; last cervical, CL; first thoracic, TF; middle thoracic, TM; last thoracic, TL; diaphragmatic vertebrae, TD; first lumbar, LF; middle lumbar, LM; last lumbar, LL.

### 2.2 Ecological categories

Carnivoran hunting type was defined based on Van Valkenburgh et al., (1985) and taxa were divided into four categories, pounce, pursuit, ambush and occasional depending on their hunting behaviour. Running speed was defined based on Galis et al., (2014), where taxa were categorised into fast, half-bound (here called intermediate) and slow based on literature and their anatomy. Generally, ambush and pursuit predators are fall into the fast category, pounce are in the intermediate category, and occasional hunters fall into the slow category. However, including both variables allow us to test for morphological differences between ambush and pursuit predators that both use fast speed but over very different durations (short and long respectively, for details on both classifications see Suppl. Table 1).

### 2.3 Landmarks

We here follow the landmarking protocol from Figueirido et al., (2021). However, we have consolidated the landmarking scheme across the whole column to 30 landmarks for each vertebra, to make each of the elements comparable and suitable for the multi element analysis. To do this, we reduced the cervical landmarks from 34 to 30 (excluding landmarks 16, 17, 33, 34), the thoracic landmarks from 32 to 30 (excluding 16, 17) and the lumbar vertebrae from 36 to 30 (excluding 16, 17, 18, 19, 35, 36). For taxa with multiple specimens, we calculated the species mean prior to phylogenetic analysis.

### 2.4 Morphological Analysis

All analyses have been performed in R version 4.0.5 (R Core Team 2021) on each subsampled vertebrae, a combination of all subsampled vertebrae for each region (cervical, thoracic and lumbar) and for the entirety of the subsampled dataset (all). Vertebrae combinations have been concatenated using ‘Morphoblocks’ (Thomas et al., 2021). Morphoblocks can be used to create morphospace for multiple elements, by putting each element into a ‘block’ which then can be used to analyse each element separately, but also all elements together as a combined consensus analysis.

Prior to any morphological analysis we applied Procrustes Superimposition to minimize the effects of size and orientation using the *gpagen()* function from the package ‘geomorph’ 4.0.3 (Adams et al., 2021). We performed a principal component analysis (PCA) for each subsampled vertebrae in ‘geomorph’, which assimilates data from all landmarks and reduces them to a set of PC scores that summarize the vertebral shape of each species. Vertebral morphology was plotted in a morphospace according to their hunting strategies (pursuit, ambush, pounce, occasional) and running speeds (fast, intermediate, slow).

We used PERMANOVA to test whether vertebral morphology is significantly different among hunting strategies and running speeds using the *pairwiseAdonis()* function in vegan 2.6–2 (Oksanen et al., 2024). This test compares the variation among groups to the variation within groups using distance matrices. To take phylogeny into account, we tested for the correlation between vertebral morphology and hunting behaviour, running speed and centroid size using phylogenetic generalized least square regression (PGLS) in the R package ‘geomorph’. For the PGLS, we used a well resolved phylogeny from Nyakatura et al., (2012). After testing single vertebrae and vertebrae regions, we identified vertebrae that show the best correlation with hunting behaviour and running speed and tested them in different combinations with one another.

To predict the ecology of the fossil taxa (running speed/hunting behaviour), we performed a canonical variate analysis (CVA), using the CVA function of the package ‘Morpho’ 2.9 (Schlager, 2017). First the PCA scores of the extant dataset were used to perform the CVA, before the fossil specimen was projected into the morphospace by using the canonical variates and matrix multiplication with the PCA eigenvectors. The Mahalanobis distances of each specimen to the mean of each ecological category was used to predict the probability of membership of each ecological category using the mahalanobis () function.

## 3 Results

### 3.1 Cervical region

Shape variation in the cervical region was examined both for individual subsampled vertebrae (CF, CM, CL; Figure 2) and combined for the whole cervical region in Morphoblocks. For the individual vertebrae, PC1 and PC2 combined explained around 50% of shape variation (CF: 51.45%, CM: 50.32%, CL: 46.76%), while for the whole region they explained 39.69%. In the individual vertebrae morphospaces, there was best differentiation on PC1 and PC2 among slow and fast carnivorans, with intermediate speed species overlapping both groups (Figure 2). Similarly, for hunting strategies, there was most overlap for occasional and pounce hunters, while pursuit and ambush hunters tend to occupy distinct regions of the morphospace (Suppl. Figure 1, 2). When examining variation across the whole cervical region, a similar pattern is seen, fast and slow runners separate whereas intermediate ones overlap in morphospace. For hunting strategies, the pounce and ambush hunters occupy separate areas of the morphospace whereas the occasional and pounce ones occupy a very large area of the morphospace (Suppl. Figure 1, 2).

**Figure 2.**
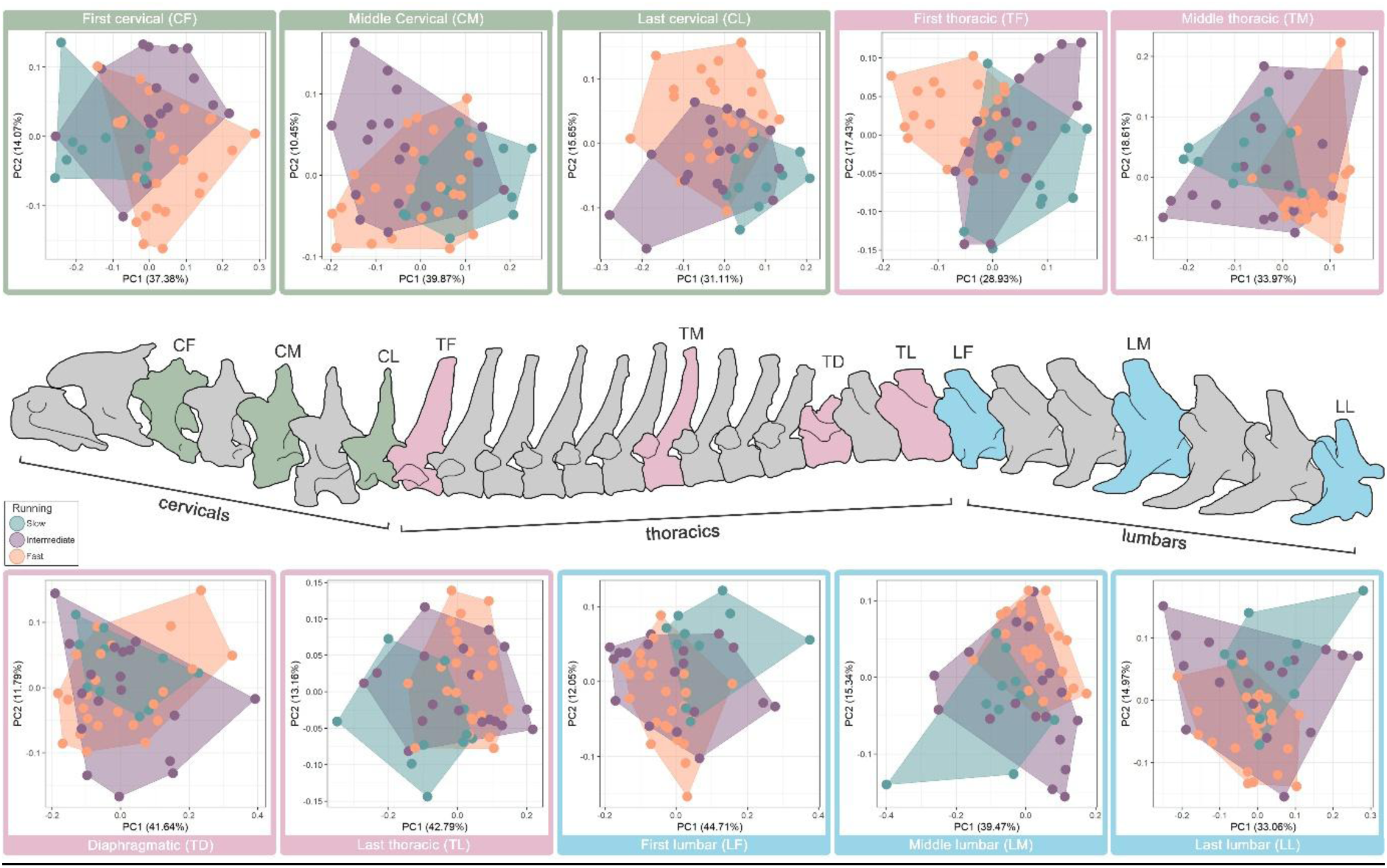
Vertebral column with marked subsampled vertebrae (green, cervicals; pink, thoracics; blue, lumbars). PCA plots show morphospace occupation of running speed (fast, intermediate, slow) for carnivoran taxa. For details on specimens and morphospace occupation for hunting strategies see Suppl. Figures 1, 3 and 5 and interactive plots: https://juliaaschwab.github.io/Vertebrae_locomotion_plots/PCA_lumbars_individual.html; https://juliaaschwab.github.io/Vertebrae_locomotion_plots/PCA_thoracics_individual.html; https://juliaaschwab.github.io/Vertebrae_locomotion_plots/PCA_lumbars_individual.html

PERMANOVA shows that cervical morphology significantly correlates with ecology for several of the speed and hunting categories (Suppl. Table 3, Suppl. Figure 7, 8). For the full cervical region, these are Fast vs. Slow (f=9.449, p=0.001), Pursuit vs. Ambush (f=8.537, p=0.001) and Ambush vs. Pounce (f=5.167, p=0.001). For single cervical vertebrae significant correlations are: Fast vs. Slow (CF: f= 9.321; CL: f= 9.370, p=0.001), Slow vs. Intermediate (CF: f= 7.292, p=0.001; CL: f= 4.676, p=0.002), Occasional vs. Ambush (CF: f= 4.742, p=0.005; CL: f= 3.485, p=0.005), Occasional vs. Pursuit (CL: f= 6.083, p=0.002), Ambush vs. Pursuit (CM: f= 12.418, p=0.001; CL: f= 6.815, p=0.001), Ambush vs. Pounce (CM: f= 5.382, p=0.003). A PGLS (Suppl. Table 7, Suppl. Figure 9) was performed to account for phylogeny and all single vertebrae were significantly correlated with running speed (CF: R2=0.175, p=0.001; CM: R2=0.172, p=0.001; CL: R2=0.162, p=0.001), as well as the full cervical region (R2=0.135, p=0.004). However, none were showed a significant correlation with hunting behaviour.

### 3.2 Thoracic region

The thoracic region was analysed as a whole, and the four subsampled vertebrae were examined separately. PC1 and PC2 explain around 50% of the overall variation for single vertebrae (TF: 46.36%, TM: 52.58%, TD: 53.43%, TL: 55.95%) and 33.86% for the thoracic region combined. Like the other regions, some of the thoracic vertebrae show better morphospace separation than others (Suppl. Figure 3, 4). TM has the best morphospace separation between slow and fast running taxa, whereas TD has the most overlap between those two running categories. Intermediate runners show a large overlap with the other groups in all four thoracic vertebrae. For TF pursuit hunters have a distinct morphospace from ambush as well as occasional hunters whereas ambush and occasional do overlap. The morphospace for TM shows a clear separation between occasional and ambush as well as occasional and pursuit, whereas ambush and pursuit occupy the same morphospace area. Both TD and TL show most overlap in those three hunting categories. The pounce category overlaps with all other groups in all four thoracic vertebrae. A similar pattern is seen for the combined thoracic region. Fast and slow runners show some separation in morphospace, whereas the intermediate category overlaps. For the hunting strategy ambush and pursuit occupy a similar area of the morphospace and separate from the occasional hunters, whereas the pounce group overlaps with all others.

Statistically significant different categories based on PERMANOVA for the thoracic region (Suppl. Table 4, Suppl. Figure 7, 8) are: Fast vs. Slow (thoracic: f= 10.681, p=0.001; TF: f= 11.108, p= 0.001; TM: f= 12.683, p= 0.001; TL: f= 7.579, p= 0.002), Fast vs. Intermediate (thoracic: f= 3.886, p=0.004; TF: f= 5.865, p= 0.001; TM: f= 6.717, p= 0.001), Occasional vs. Ambush (thoracic: f= 9.256, p=0.001; TF: f= 6.525, p= 0.001; TM: f= 12.620, p= 0.001; TL: f= 6.744, p= 0.005), Occasional vs. Pursuit (thoracic: f= 9.931, p=0.001; TF: f= 12.404, p= 0.001; TM: f= 11.329, p= 0.001), Ambush vs. Pursuit (TF: f= 9.315, p= 0.001; TL: f= 4.572, p= 0.003), Ambush vs. Pounce (TM: f= 4.306, p= 0.002) and Pursuit vs. Pounce (TF: f= 4.569, p= 0.002). By contrast, for the phylogenetically-corrected analysis (PGLS), the first and last thoracic vertebrae, as well as all thoracic combined (thoracic: R2=0.1669, p=0.001; TF: R2=0.179, p=0.001; TL: R2=0.172, p=0.002), were significantly correlated with running speed (Suppl. Table 8, Suppl. Figure 9). There were no significant correlations with hunting behaviour.

### 3.3 Lumbar region

Variation in shape was analysed for the entire lumbar region and the three subsampled vertebrae were analysed separately. PC1 and PC2 of the full lumbar region shows shape variation of 45.55% and the individual lumbar vertebrae around 50% (LF: 56.76%, LM: 54.34%, LL: 48.03%). For the lumbar region, PCA analysis and morphospace occupation is comparable to other regions, with LF generally showing the best separation between different groups. Fast and slow show some separation in all lumbars, as well as in the combined lumbar vertebrae, whereas the intermediate category overlaps with all of them (Suppl. Figure 5, 6). For the hunting strategies, ambush and pursuit show some overlap of morphospace area, but separation from occasional hunters (Suppl. Figure 5, 6). The pounce category overlaps with all others and has no clear separation, for single lumbars as well as the full lumbar region combined.

Lumbar vertebrae show significant correlations (based on PERMANOVA) with Fast vs. Slow (lumbar: f= 9.931, p=0.001; LF: f= 9.600, p= 0.001; LM: f= 7.881, p= 0.001; LL: f= 5.239, p= 0.001), Occasional vs. Ambush (lumbar: f= 9.264, p=0.001; LF: f= 8.139, p= 0.002; LM: f= 7.415, p= 0.001; LL: f= 5.969, p= 0.002) and Ambush vs. Pursuit (LF: f= 5.758, p= 0.001) (Suppl. Table 5, Suppl. Figure 7, 8). In the phylogenetically corrected analysis (Suppl. Table 9, Suppl. Figure 9), all single lumbar vertebrae, as well as total lumbar morphology combined, correlated significantly with running speed (lumbar: R2=0.161, p=0.001; LF: R2=0.182, p=0.001; LM: R2=0.180, p=0.001; LL: R2=0.149, p=0.002). LF was the only subsampled vertebrae that was significantly correlated with hunting behaviour (LF: R2=0.172, p=0.005).

### 3.4 Overall vertebral column shape

To examine overall variation in shape across the whole column, the ten subsampled vertebrae were placed into ‘blocks’ which then can be analysed as multiple-part objects using Morphoblocks. Principal component analysis (PCA) was performed to show variation across carnivorans associated with running speed as well as hunting behaviour (Figure 3). The first two principal components explained around 19% and 12% of variation respectively. PC1 represents variation from a more elongate vertebral column in with strongly inclined lumbar transverse processes at positive scores, to a more compressed vertebral column, with taller and straighter neural spines at negative scores (Figure 3). PC2 represents variation from shorter and more robust neural spines at positive scores, to narrower, more elongate and curved neural spines at negative scores. Generally, taxa showing ambush and pursuit hunting behaviour are constrained to the positive PC1 region of morphospace, whereas the occasional and pounce ones are more widespread. Fast runners tend to have more positive PC1 scores and negative PC2 scores than slow runners, while intermediate runners are broadly distributed.

**Figure 3.**
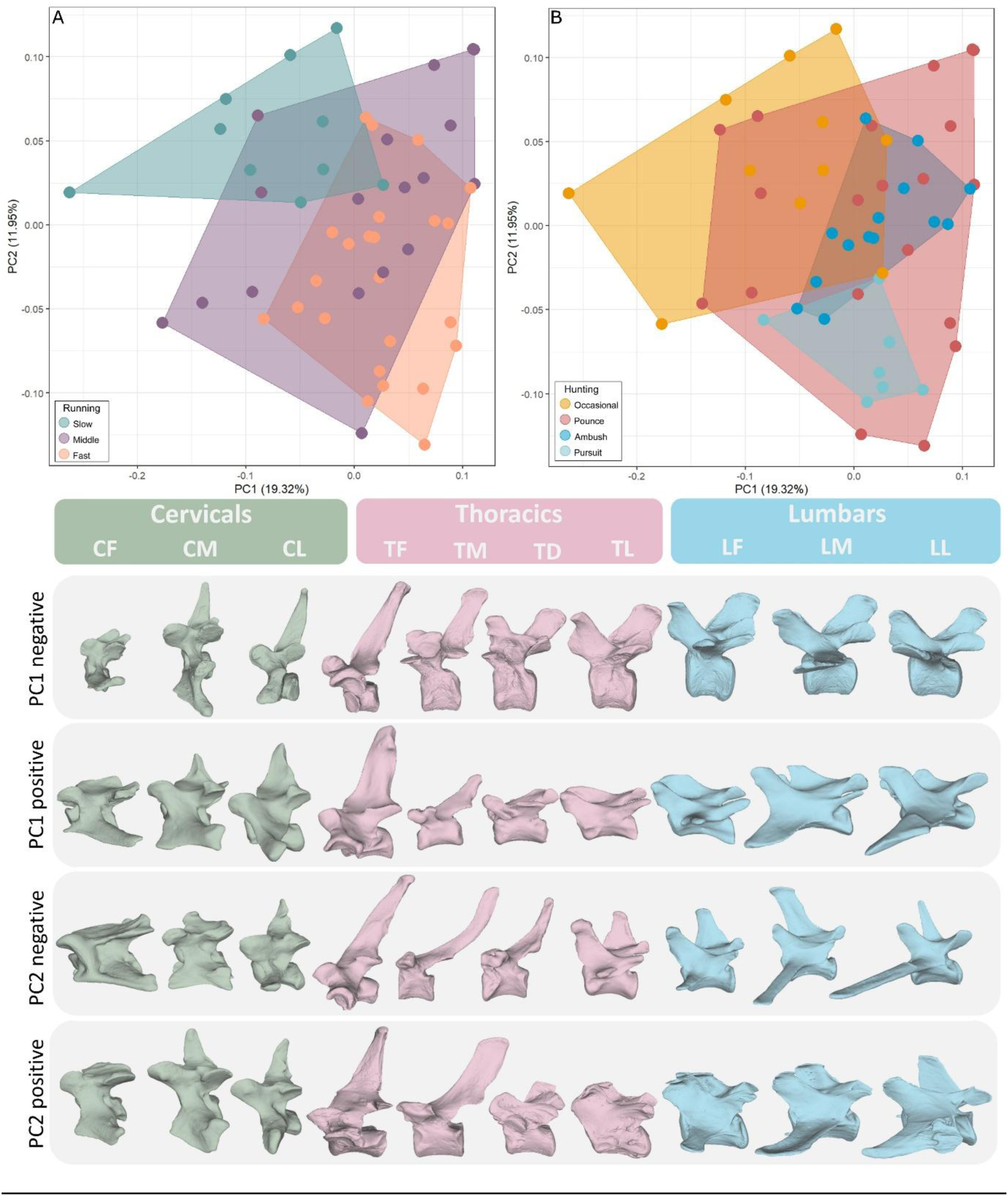
Vertebral shape morphospaces, showing the distribution extant carnivoran taxa of different running speeds (A) and hunting behaviour (B), based on principal component analysis of 3D landmarks. Based on all ten subsampled vertebrae combined using morphoblocks. Vertebral shapes are represented by PC1 negative *Ailuropoda melanoleuca*; PC1 positive *Mustela putorius*; PC2 negative *Otocyon megalotis*; PC2 positive *Ailurus fulgens*. For details on taxa see the interactive plot here: https://juliaaschwab.github.io/Vertebrae_locomotion_plots/PCA_full_column.html

The morphospace occupation test (PERMANOVA) supports the observation that some hunting behaviours and running speeds are significantly different (p-value ≤ 0.005) from each other (Suppl. Table 6, Suppl. Figure 7, 8). Significant combinations were Fast vs. Slow (f=11.803, p=0.001), Fast vs. Intermediate (f=4.209, p=0.003), Occasional vs. Ambush (f= 8.582, p=0.001), Occasional vs. Pursuit (f= 6.958, p=0.001), Ambush vs. Pursuit (f= 6.201, p=0.001) and Ambush vs Pounce (f= 3.940, p=0.004). However, the correlation between vertebrate morphology and ecological parameters might be partially influenced by phylogeny. Therefore, we also test the relationship between vertebral shape and running speed, hunting behaviour and centroid size using PGLS (Figure 4; see Suppl. Table 10 and Suppl. Figure 10 for full results). Overall vertebral column shape shows a significant correlation with running speed (R2= 0.135, p=0.004), but not with hunting behaviour.

**Figure 4.**
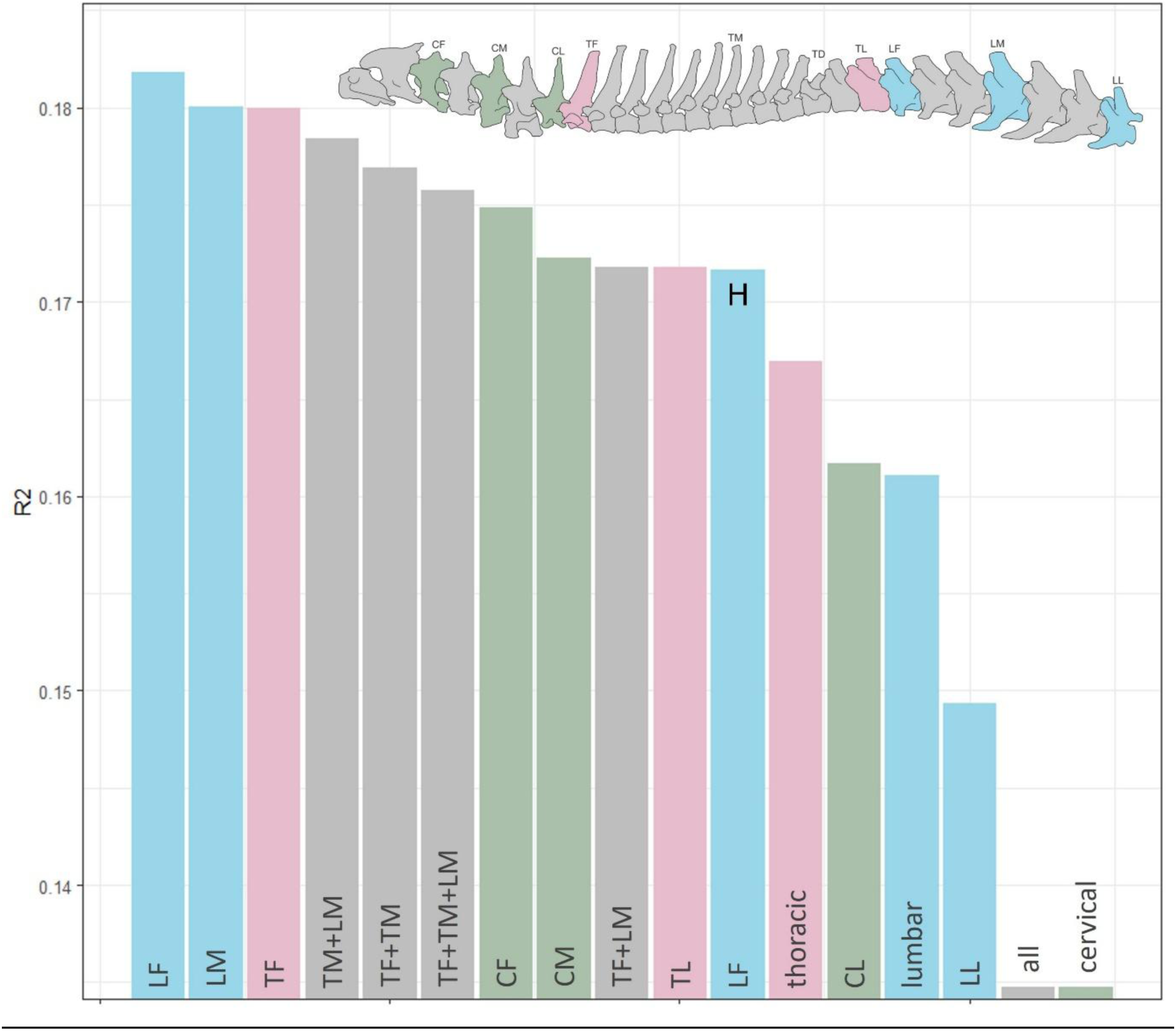
Plot of PGLS R^2^ values of running speed and hunting behaviour (marked with an H). Only significant values (p-value ≤ 0.005) are shown.

### 3.5 Vertebrae combinations

We tested if correlation between vertebral morphology and running behaviour and/or hunting strategy can be improved with different combinations of individual vertebrae using PGLS (see Suppl. Table 10 for details). There were significant correlations between vertebral morphology and running speed for all the combinations of vertebrae tested, so the effect size of the analyses (r-squared) was compared to examine variations in goodness of fit. The strongest correlations of vertebral morphology are seen with running behaviour (R2 > 0.153). Further, some vertebrae that did not have significant relationships on their own, became significant when paired with other vertebrae, and so adding more vertebrae can add more information to an analysis. However, adding more vertebrae did not always improve the model fit. Surprisingly, the overall effect size of running speed was strongest for the isolated vertebrae TF, LF and LM, and still high for CF, CM and TL indicating that individual vertebrae can actually be better predictors of ecology than multiple vertebrae (Figure 4). The vertebral combinations with the best relationship were TM+LM, TF+TM, TF+TM+LM and TF+LM. The combined thoracic and lumbar regions also had a good fit, with R2 around 16%, but the cervical region alone, and the total overall column shape had much worse fit (R2=0.135).

### 3.6 CVA and Fossil predictions

Based on the results of the PGLS, we selected 6 vertebrae that had the strongest relationship with the ecological variables (R^2^ over 0.16) to predict running speed and hunting ecology in the extinct species, *Canis dirus*. For running speed, these were CF, CM, CL, TL, LF and LM, while for hunting behaviour this was only LF. Some vertebrae were not analysed due to low R^2^ scores (LL), non-significance during PGLS (TD, TM), or because there were no available fossil specimens with good enough preservation (TF, TM). Further it was not possible to test for the vertebrae combinations that showed the highest correlation with running speed, as TM was not preserved in *C. dirus*. However, a combination of all preserved vertebrae was tested.

We performed a canonical variate analysis (CVA) of the PC scores for these vertebrae to test if ecological categories can be discriminated based on their morphology (Figure 5). The classification accuracy for running speed was highest in CF (77%) and 58% on average, which is significantly higher than for hunting behaviour (37%). The highest r-squared values from PGLS were from the first lumbar, which has an overall classification rate of 60.47% for running speed (see Suppl. Table 11) and 37.21% for hunting behaviour (see Suppl. Table 12). Next, landmarks taken from specimens of *Canis dirus*, whose ecology is unknown, were projected back into the CVA space for CF, CM, CL, TL, LF and LM. Examining the CVA plot for the first lumbar (highest R^2^ from PGLS, Figure 5), each running speed occupies a distinct portion of morphospace, while for hunting behaviour, only pursuit predators are clearly distinguished. The fossil specimen *Canis dirus* plots closest to the fast and pursuit morphospaces. Similarly, in the CVAs for CF and TL (Suppl. Figure 11, 14), the fossil specimen plots near the fast morphospace, while for LM (Suppl. Figure 15) it falls closest to the intermediate region. For CM and CL (Suppl. Figure 12, 13), it even plots within the intermediate morphospace. When analysing a combined CVA of all preserved vertebrae (Suppl. Table 13; Suppl. Figure 16), there is considerable overlap between ecological categories, with *Canis dirus* falling within both fast and slow morphospaces.

**Figure 5.**
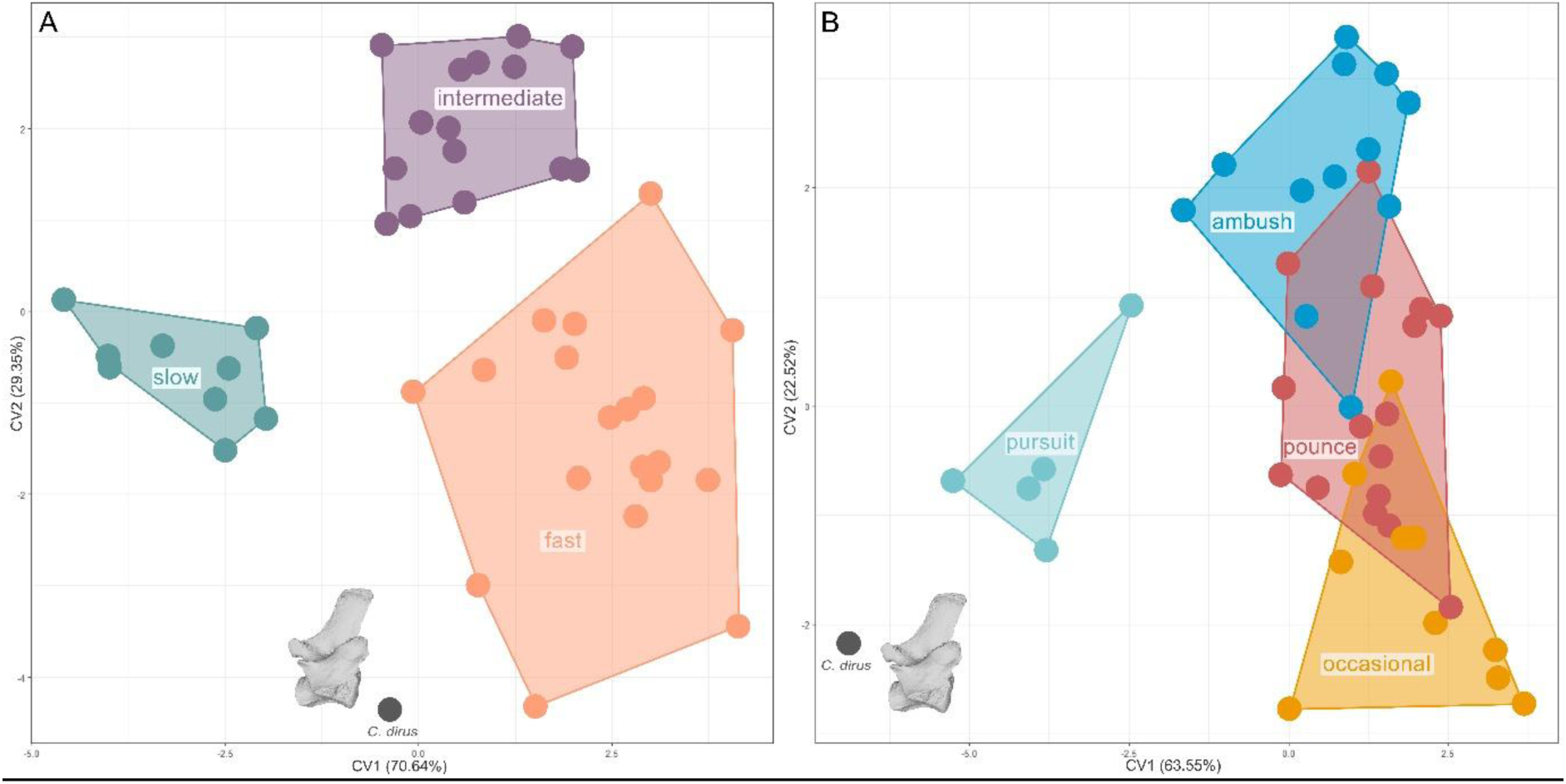
Canonical variate analysis (CVA) based on LF, showing shape morphospace, the distribution of extant carnivorans and projected *Canis dirus* in a canonical variate analysis morphospace based on running speed (A) and hunting behaviour (B). For details on taxa see the interactive plot here: https://juliaaschwab.github.io/Vertebrae_locomotion_plots/CVA_LF_running.html; https://juliaaschwab.github.io/Vertebrae_locomotion_plots/CVA_LF_hunting.html

To statistically test the likelihood of group membership for the fossil taxa, we estimated the probability for *C. dirus* and its vertebrae using Mahalanobis distances (Table 1). For running speed, CF, CM, CL, TL, LF and LM were used as they show the highest correlation (highest R^2^ values in PGLS). Strongly supported predictions were classified as those with a probability greater than 0.95 (Suppl. Table 15). *Canis dirus* was recovered as a fast runner with strong support in three out of six metrics (CF: 0.99, TL: 0.99, LF: 0.99) and as an intermediate runner with strong support in two (CM:0.99, CL:0.99). LM does not show a good prediction with the intermediate category (prob.= 0.57), as well as the combination of all vertebrae for fast (prob.= 0.79). For hunting behaviour, we only used LF as this is the only vertebrae that has been significantly correlated, where *Canis dirus* (prob.= 0.99) is predicted to be a pursuit hunter (Suppl. Table 14).

**Table 1.** Running speed predictions for the fossil *Canis dirus* using Mahalanobis distances.

| Vertebra | Category | Prediction | Probability |
| --- | --- | --- | --- |
| CF | Running | fast | 0.999 |
| CM | Running | intermediate | 0.992 |
| CL | Running | intermediate | 0.999 |
| TL | Running | fast | 0.999 |
| LF | Running | fast | 0.999 |
| LF | Hunting | pursuit | 0.999 |
| LM | Running | intermediate | 0.573 |
| all | Running | fast | 0.795 |

## 4 Discussion

Ecomorphological studies in carnivorans have been a major research focus because of their broad ecological range and role as keystone predators in many ecosystems (for an overview see Schwab et al. 2023). The vertebral column is an important component of the locomotory system that can provide insights into the ecology and evolution of extinct carnivorans (Álvarez et al. 2013; Jones 2015; Randau et al. 2016; Esteban et al. 2020; Figueirido et al. 2021; Martín-Serra et al. 2021). However, the relationship between the morphology of the vertebral column and ecological traits has rarely been used to predict behaviour for fossil taxa. A vast majority of ecomorphological research focuses on single elements or ratios between elements, such as limb morphology, which has been the focus of previous locomotor evolution studies (Samuels et al. 2013). Different aspects of limb morphology have been linked with hunting behaviour in carnivorans (Harris et al. 1997; Iwaniuk et al. 1999; Andersson et al. 2004; Meachen-Samuels et al. 2009a, b; Figueirido et al. 2015; Figueirido et al. 2023), but also with habitat (Gonyea et al. 1978; Figueirido et al. 2016). However new methods can help to overcome the challenges of studying multiple elements and produce a better holistic picture of ecomorphological signals. Here we use ‘Morphoblocks’ (Thomas et al. 2021) to examine the variation across multiple serial elements and compare combinations with isolated vertebrae. Using 3D geometric morphometrics and a multi-element approach, we have examined the ecological signal of locomotor mode and running speed in carnivoran vertebrae and predicted ecology in fossil taxa using their vertebral morphology. We show that vertebral morphology correlates most strongly with running speed, and that isolated vertebrae can provide as much ecological signal as combinations of multiple elements.

### 4.1 Vertebral morphology correlates with running speed in carnivorans

Vertebral morphology correlates more strongly with running speed than with hunting mode, a pattern consistent across all three regions of the vertebral column. Principal component analysis reveals distinct patterns of morphospace occupation corresponding to different running speeds among carnivorans (Figure 3). Fast runners, typically ambush and pursuit predators such as felids and canids, occupy tightly clustered regions of morphospace, indicating specialised vertebral adaptation for rapid, powerful locomotion essential to their speed and agility. In contrast, slower exhibit more dispersed morphospaces, indicating generalized and varied locomotor adaptations, with intermediate-speed taxa positioned between these two extremes. Certain vertebral regions show stronger correlations with running speed than others. The first and middle lumbar vertebrae, along with the first thoracic vertebra, show the strongest association with running speed (Figure 4) and are thus the most reliable for inferring locomotion in fossil taxa. Significant differentiation between fast and slow runners is evident across all vertebral regions (cervical, thoracic, lumbar, and combined), as demonstrated by PERMANOVA results. These findings support the hypothesis that vertebral morphology is a robust predictor of running speed in carnivorans.

The morphological variation related to running speed include the anterior–posterior length of the vertebral centrum and the size and shape of the neural spine (Figure 3). On the negative end of PC1 (slower speed), vertebrae tend to have narrower centra and taller, straighter, and narrower neural spines, features associated with increased spinal stability and muscle attachment. For PC2, variation centres on the width of the neural spine and centrum length, though these differences are less pronounced. On the negative side of PC2 (faster speed), centra are wider antero-posteriorly, and neural spines are longer, thinner, and more curved, likely reflecting a trade-off between vertebral mobility and stability,

Similar patterns of vertebral variation have been observed in other mammals (Kort and Polly, 2023). Vertebral centrum and disk shape correlate with the spine’s range of motion, especially in the lumbar region, by influencing flexion, extension, and lateral movement between adjacent vertebrae (Jones et al., 2021). It has been demonstrated that longer, more elongated centra permit greater spinal flexibility, which enhances stride length and mobility critical for fast-moving species. In contrast, shorter or more robust centra can restrict motion, contributing to greater spinal stability. Neural spines primarily relate to muscle attachment, a taller spine increases surface area and mechanical advantage, enhancing force transmission and spinal stabilization (Granatosky et al., 2014). Fast-running carnivorans commonly exhibit more elongated and slender vertebrae with longer neural spines and transverse processes, enhancing spinal flexion and extension to increase stride length and providing larger moment arms for muscle attachment. These adaptations facilitate efficient force generation and stabilise the spine during rapid locomotion.

These morphological distinctions enable reliable inferences about running speed in carnivorans. Lumbar vertebral shape shows strong correlations with locomotor habits (Boszczyk et al., 2001; Alvarez et al., 2013; Figueirido et al., 2021), with the degree of vertebral movement linked to centrum size, elongated centra enable greater flexibility (Long et al., 1997; Pierce, Clack and Hutchinson, 2011). Our findings align with this pattern, as faster, more active carnivorans tend to have more elongated vertebrae, consistent with the demands of their locomotor strategies. By contrast, differences in hunting mode are often more strongly reflected in the appendicular skeleton, particularly the forelimbs directly involved in prey capture and handling. Ambush predators have robust forelimbs for grappling prey, while pursuit predators often show more gracile, elongated limbs adapted for speed and endurance (Meachen-Samuels and Van Valkenburgh, 2009a, b). Because hunting strategies primarily influence how prey is captured, rather than how it is approached, selective pressures on the axial skeleton are less consistent. This distinction helps explain why vertebral morphology better predicts running speed than predatory strategy.

This pattern supports the concept that different ecological behaviours are reflected in distinct skeletal regions. While the axial skeleton, especially the lumbar vertebrae, can predict running speed effectively, other ecological traits such as predatory strategy or habitat use may be better inferred from cranial or appendicular anatomy. This underscores the value of a multi-element approach in ecomorphological studies, analysing multiple anatomical regions to capture the full spectrum of ecological adaptations. Trade-offs among functional traits across skull, appendicular, and axial skeletons (Law et al., 2025) demonstrate that varied ecological demands differentially shape body regions. Therefore, understanding morphology–ecology relationships in an evolutionary context requires integrating data across the entire skeleton.

### 4.2 Isolated vertebrae can provide as much ecological signal as multiple vertebrae or whole regions

Here, we compare the ecological signal of isolated vertebrae to that obtained by analysing multiple vertebrae or entire regions, to evaluate the utility of isolated vertebrae for interpreting locomotor ecology in the fossil record. Most single vertebrae maintained significant correlations with running speed. For instance, the first lumbar vertebra (LF) showed robust correlations with both hunting behaviour and running speed (Figure 4), even when phylogenetic effects were considered, suggesting this vertebra may have evolved specific adaptations critical for locomotion across different lineages. The three strongest effects were found in isolated vertebrae, the first lumbar (LF), middle lumbar (LM), and first thoracic (TF), although certain vertebral combinations also showed meaningful associations. This indicates that although multi-vertebral analyses can be beneficial, they are certainly not essential to extract ecological signal from vertebral morphology.

The lumbar region is key to sagittal flexion and extension of the spine during running (Figueirido et al., 2021; Belyaev et al., 2024). This region is highly mobile and acts as a dynamic bridge between the thorax and pelvis, contributing significantly to stride length and body propulsion. Elongated lumbar vertebrae with long neural spines support both enhanced flexibility and the muscle attachment needed for dorsoventral flexion, particularly in fast, agile species (Pierce et al., 2011; Belyaev et al., 2024). The high correlation seen in the first lumbar vertebra (LF) may reflect its critical role in facilitating sagittal bending of the spine because the mid-spine region, constituting the post diaphragmatic thoracic vertebrae and the anterior lumbar vertebrae, contribute most to sagittal bending during running. Interestingly, although the diaphragmatic thoracic vertebra (TD) lies within this biomechanically important transitional zone, it did not show a significant correlation with running speed. This suggests that not all vertebrae in the contribute equally to locomotor performance. The first thoracic vertebra (TF) likely plays an important role. It marks the transition from the cervical to thoracic region and may act as an anchor point for both neck and trunk musculature. Its positioning suggests it may be involved in transferring force between the forelimbs and the axial skeleton during high-speed locomotion (Schilling and Hackert 2006). Moreover, variation in the shape and curvature of the thoracic neural spines, as seen along PC2, might reflect differing demands for stability versus flexibility among taxa with distinct locomotor strategies (Randau and Goswami, 2017). Surprisingly, the cervical region, also showed relatively high predictive power despite being less directly involved in locomotor movements of the limbs. Cervical vertebrae control head movement and orientation, which are essential during hunting, particularly in ambush predators that rely on rapid, precise head and neck strikes (Belyaev et al., 2024).

One explanation for why isolated vertebrae, such as LF, LM, and TF, often provide a clearer ecological signal than multiple vertebrae or entire regions combined is that different vertebrae reflect distinct functional and ecological adaptations. Vertebrae from different axial regions experience region specific mechanical demands and thus respond to ecological pressure in different ways. For example, lumbar vertebrae primarily facilitate spinal flexibility and propulsion (Pierce et al., 2012; Slijper, 1946), while thoracic vertebrae contribute more to ribcage stabilization and postural support (Granatosky et al., 2014; Taylor et al., 2015). As a result, when vertebrae from multiple regions are combined into a single analysis, the resulting morphological signal may average or blur these region-specific adaptations. This blending of signals can obscure the more precise ecological information captured by isolated, functionally distinct vertebrae (Jones et al., 2021; Figueirido et al., 2021).

Additionally, vertebrae are subject to different selective pressures based on their location and function within the axial skeleton. For instance, the first lumbar vertebra frequently shows morphological specializations related to muscle attachment and spinal flexion, supporting rapid acceleration and agility in cursorial mammals (Boszczyk et al., 2001; Slijper, 1946). In contrast, cervical vertebrae morphology is more influenced by head mobility and feeding behaviours (Taylor et al., 2015). These region-specific adaptations highlight why isolated vertebrae can serve as powerful proxies for ecological inference, particularly in paleontological contexts where complete vertebral series are rarely preserved. Their ability to retain strong ecological signals, even in isolation, underscores their value for reconstructing locomotor behaviour and predatory strategies in fossil taxa (Figueirido et al., 2021; Jones et al., 2021). Indeed, we show with the example provided below, that analysing all available elements together, which may be considered the default approach, reduces the predictive power of the ecological analysis. Therefore, we advocate for palaeontologists to conduct analyses separately on different functional regions of the spine, and predict that the ecological signals may be mixed, requiring more nuanced functional and ecological interpretations based on mosaic ecological patterns across the vertebral column.

### 4.3 Canis dirus was likely a fast, pursuit predator

The vertebral morphology of *Canis dirus* suggest that, based on Mahalanobis distance analyses (Anyonge and Baker, 2006), it was a fast, active predator with a pursuit hunting style (Table 1) comparable to that of modern the modern wolf, *Canis lupus*. In the CVA, *C. dirus* plots close to, but just outside, of the modern morphospace, suggesting a unique morphology and locomotor style not directly comparable to any extant species. While it shares many ecological and morphological traits with modern wolves, *C. dirus* differed notably in its cranial anatomy, possessing broader, more robust skulls, deeper jaws, and significantly larger, more massive teeth (Merriam, 1912). These features have led to contrasting interpretations of its feeding behaviour. Some studies propose that *C. dirus* engaged in bone-crushing feeding strategies like those of extant spotted hyenas (*Crocuta crocuta*) or extinct Miocene borophagine dogs (Kurtén and Anderson, 1980; Van Valkenburgh and Ruff, 1987; Biknevicius and Ruff, 1992; Biknevicius and Van Valkenburgh, 1996; Meehan and Martin, 2003). Supporting this interpretation, its canines are morphologically more similar to those of bone-crushing carnivores than to modern canids (Van Valkenburgh and Ruff, 1987). However, other evidence, such as relative jaw strength, points toward a predatory and feeding behaviour more akin to that of modern wolves, indicating a potential mix of adaptations (Biknevicius and Van Valkenburgh, 1996) for both scavenging and pursuit predation (Figueirido et al. 2015). These conflicting interpretations reflect ongoing debate about whether *C. dirus* was a specialized bone-crusher or primarily a flesh-feeding predator with some bone-processing capabilities.

Although the vertebral morphology of *C. dirus* suggests locomotor adaptations for speed, the classification across different vertebrae is not entirely consistent. Posterior thoracic and lumbar elements, associated with sagittal bending during running, as well as the first cervical, strongly suggest a fast-running speed for *C. dirus.* In contrast, the other cervicals, which are more involved in head movement and control, align more closely with intermediate taxa. This likely reflects the biological and functional complexity of the vertebral column, where different regions serve distinct roles. Lumbar vertebrae drive flexion-extension and stride length, thoracic vertebrae stabilize the trunk and transmit force, and cervical vertebrae enable rapid head movements crucial essential for prey capture. As such, individual vertebrae may evolve semi-independently in response to different selective pressures, predation style, prey size, terrain, or social hunting behaviour, leading to a mosaic of signals. This region-specific variation is consistent with findings that serial disparity along the carnivoran vertebral column reflects complex, regionally differentiated adaptive roles shaped by metameric evolution (Figueirido et al. 2021). For instance, the robust cranial features and potential bone-crushing behaviour of *C. dirus* may require increased robustness of the cervical vertebrae, potentially overlaying the ecological signal for high-speed locomotion, and resulting in a more ‘intermediate speed’ morphology. These regionally variable signals may therefore reflect a combination of behavioural traits not well captured by single ecological categories. Moreover, there are inherent limitations to applying categorical classification to continually varying functional traits such as running speed. Locomotor and predatory strategies in carnivorans exist along ecological gradients rather than in discrete bins. Consequently, classification uncertainty or overlap between fast and intermediate running speed morphospaces may indicate ecological variability in extinct species, as are commonly observed among living populations. For extinct taxa such as *C. dirus*, this pattern may reflect a locomotor strategy that combined endurance-based pursuit with opportunistic scavenging, a unique combination not observed among the extant fauna. These mixed signals reinforce with the view that *C. dirus* was not simply a Pleistocene analogue of the modern wolf but a distinctive predator with its own adaptive niche.

These findings underscore the power of vertebral morphology in reconstructing the locomotor and ecological adaptations of extinct carnivorans. Even when overall classification accuracy is moderate, certain vertebrae, particularly those with key biomechanical roles like the first lumbar (LF), middle lumbar (LM), and first thoracic (TF), retain strong, consistent ecological signals. This suggests that isolated elements can capture highly specific functional adaptations that may become diluted when multiple vertebrae with differing roles are analysed together. Rather than requiring full sequences, meaningful ecological inference can be drawn from single, strategically informative elements. As such, vertebral morphology offers a robust and flexible tool for interpreting locomotor behaviour and ecological strategies in fossil taxa, even when skeletal remains are incomplete. Further, vertebrae are particularly effective at reflecting aspects of locomotor performance such as speed and spinal flexibility because they directly influence stride length and force transmission along the body axis. However, they are less informative about specific hunting modes or behaviour specialisations. In contrast, limb morphology provides clearer insights into locomotor mode and substrate use, as limbs mediate direct interaction with the environment, though limb shape may correlate less directly with speed due to their multifunctional roles beyond locomotion. Together, these complementary anatomical perspectives deepen our understanding of extinct carnivoran ecology, emphasizing the value of integrating axial and appendicular data in palaeobiological reconstructions.

## Author contributions

JAS designed and performed the research, analysed the data and wrote the paper; KEJ provided funding, helped with the interpretation of the data and reviewed drafts of the paper; BF provided data and reviewed drafts of the paper.

## Acknowledgement

We thank Zena Timmons (NMS), Tiago Metello (NMS) and William Simpson (FMNH) for providing generous access to the specimens in their care. We are also grateful to Logan Anders who helped with data collection. This work was supported by the Royal Society (RF\ERE\210253 to K.E.J.) and a Leverhulme Trust Early Career Fellowship (ECF-2023-365 to J.A.S.).

