## Supplements for "Ecological inference from isolated vertebrae: Evaluating functional signal across the carnivoran spine"

### 1. Specimen List

Suppl. Table 1. Detailed information on the scanned specimens; their museum specimen numbers, source and assigned hunting behaviour/running speeds.

| Taxa | Family | ID | Source | Hunting behaviour | Running speed |
|---|---|---|---|---|---|
| ***Ailurus fulgens*** | **Ailuridae** | UVA 1802 | Figueirido et al. 2021 | Occasional | slow |
| ***Canis lupus*** | **Canidae** | UVA 4589 | Figueirido et al. 2021 | Pursuit | fast |
| ***Chrysocyon brachyurus*** | **Canidae** | NMS Z.2015.177 | Figueirido et al. 2021 | Pounce | fast |
| ***Cuon alpinus*** | **Canidae** | UVA 7106 | Figueirido et al. 2021 | Pursuit | fast |
| ***Lycaon pictus*** | **Canidae** | UVA 1167 | Figueirido et al. 2021 | Pursuit | fast |
| ***Speothos venaticus*** | **Canidae** | NMS Z.2015.96.4 | Figueirido et al. 2021 | Pounce | intermediate |
| ***Vulpes vulpes*** | **Canidae** | UVA 150 | Figueirido et al. 2021 | Pounce | fast |
| ***Otocyon megalotis*** | **Canidae** | NMS PH107.04 | Scanned by JAS at the National Museum of Scotland, Edinburgh | Pounce | intermediate |
| ***Vulpes lagopus*** | **Canidae** | NMS A.F.I. | Scanned by JAS at the National Museum of Scotland, Edinburgh | Pounce | intermediate |
| ***Canis lupus*** | **Canidae** | MU 073 | Scanned by Logan Anders, University of Manchester | Pursuit | fast |
| ***Canis lupus*** | **Canidae** | MU 071 | Scanned by Logan Anders, University of Manchester | Pursuit | fast |
| ***Vulpes vulpes*** | **Canidae** | MU 091a | Scanned by Logan Anders, University of Manchester | Pounce | fast |
| ***Cryptoprocta ferox*** | **Eupleridae** | NMS EA36.08 | Figueirido et al. 2021 | Pounce | fast |
| ***Acinonyx jubatus*** | **Felidae** | UVA, NN | Figueirido et al. 2021 | Pursuit | fast |
| ***Caracal caracal*** | **Felidae** | UVA 1556 | Figueirido et al. 2021 | Ambush | fast |
| ***Felis silvestris*** | **Felidae** | UVA NN | Figueirido et al. 2021 | Ambush | fast |
| ***Leptailurus serval*** | **Felidae** | UVA 6070 | Figueirido et al. 2021 | Ambush | fast |
| ***Lynx canadiensis*** | **Felidae** | UVA Z.2001.117.9 | Figueirido et al. 2021 | Ambush | fast |
| ***Neofelis nebulosa*** | **Felidae** | NMS Z.2001.8 | Figueirido et al. 2021 | Ambush | fast |
| ***Panthera leo*** | **Felidae** | UVA NN | Figueirido et al. 2021 | Ambush | fast |
| ***Panthera pardus*** | **Felidae** | UVA 1600 | Figueirido et al. 2021 | Ambush | fast |

| | | | | | |
|---|---|---|---|---|---|
| ***Panthera tigris*** | **Felidae** | MNCN 21574 | Figueirido et al. 2021 | Ambush | fast |
| ***Panthera tigris altaica*** | **Felidae** | | Figueirido et al. 2021 | Ambush | fast |
| ***Puma concolor*** | **Felidae** | UVA 409 | Figueirido et al. 2021 | Ambush | fast |
| ***Lynx rufus*** | **Felidae** | NMS R6800 | Scanned by JAS at the National Museum of Scotland, Edinburgh | Ambush | interm ediate |
| ***Panthera onca*** | **Felidae** | NMS Z.2013.162 | Scanned by JAS at the National Museum of Scotland, Edinburgh | Ambush | fast |
| ***Panthera uncia*** | **Felidae** | NMS Z.2009.024.001 | Scanned by JAS at the National Museum of Scotland, Edinburgh | Ambush | fast |
| ***Panthera leo*** | **Felidae** | MU 127 | Scanned by Logan Anders, University of Manchester | Ambush | fast |
| ***Cynictis penicillata*** | **Herpestidae** | UVA 6730 | Figueirido et al. 2021 | Pounce | slow |
| ***Herpestes ichneumon*** | **Herpestidae** | UVA 7344 | Figueirido et al. 2021 | Pounce | slow |
| ***Suricatta suricatta*** | **Herpestidae** | UVA 3098 | Figueirido et al. 2021 | Pounce | interm ediate |
| ***Suricatta suricatta*** | **Herpestidae** | UVA 7103 | Figueirido et al. 2021 | Pounce | interm ediate |
| ***Crocuta crocuta*** | **Hyaenidae** | UVA 4215 | Figueirido et al. 2021 | Pursuit | fast |
| ***Hyaena hyaena*** | **Hyaenidae** | UVA 2981 | Figueirido et al. 2021 | Pounce | interm ediate |
| ***Proteles cristatus*** | **Hyaenidae** | NMS PH37.98 | Figueirido et al. 2021 | Occasion al | interm ediate |
| ***Martes foina*** | **Mustelidae** | UVA 7342 | Figueirido et al. 2021 | Pounce | interm ediate |
| ***Mustela putorius*** | **Mustelidae** | UVA 7203 | Figueirido et al. 2021 | Pounce | interm ediate |
| ***Mustela putorius*** | **Mustelidae** | UVA 7343 | Figueirido et al. 2021 | Pounce | interm ediate |
| ***Martes foina*** | **Mustelidae** | NMS R21399 | Scanned by JAS at the National Museum of Scotland, Edinburgh | Pounce | interm ediate |
| ***Meles meles*** | **Mustelidae** | MU 056 | Scanned by Logan Anders, University of Manchester | Occasion al | slow |
| ***Meles meles*** | **Mustelidae** | MU 125a | Scanned by Logan Anders, University of Manchester | Occasion al | slow |
| ***Nasua narica*** | **Procyonidae** | UVA 5542 | Figueirido et al. 2021 | Occasion al | slow |
| ***Potos flavus*** | **Procyonidae** | UVA 124 | Figueirido et al. 2021 | Occasion al | slow |

| | | | | | |
|---|---|---|---|---|---|
| ***Procyon lotor*** | **Procyonidae** | UVA 4264 | Figueirido et al. 2021 | Occasion al | slow |
| ***Nasua nasua*** | **Procyonidae** | NMSZ.2015.163.2 | Scanned by JAS at the National Museum of Scotland, Edinburgh | Pounce | interm ediate |
| ***Ailuropoda melanoleuca*** | **Ursidae** | NMS Z.1986.19 | Figueirido et al. 2021 | Occasion al | slow |
| ***Ursus arctos*** | **Ursidae** | NMS Z.2003.41.1 | Figueirido et al. 2021 | Occasion al | interm ediate |
| ***Ursus maritimus*** | **Ursidae** | MNCN 21570 | Figueirido et al. 2021 | Pounce | interm ediate |
| ***Arctictis binturong*** | **Viverridae** | UVA 5474 | Figueirido et al. 2021 | Occasion al | slow |
| ***Genetta genetta*** | **Viverridae** | UVA 1488 | Figueirido et al. 2021 | Pounce | interm ediate |
| ***Genetta tigrina*** | **Viverridae** | UVA 3824 | Figueirido et al. 2021 | Pounce | interm ediate |
| ***Paradoxurus hermaphrodi tus*** | **Viverridae** | UVA 4610 | Figueirido et al. 2021 | Occasion al | interm ediate |

Suppl. Table 2. Detailed information on the scanned fossil specimen; their museum specimen numbers, source and elements.

| **Taxa** | **Family** | **Element** | **ID** | **Source** |
|---|---|---|---|---|
| ***Canis dirus*** | Canidae | CF | ROM 27396 | Scanned by JAS at the Royal Ontario Museum, Toronto |
| | | CM | ROM 27460 | |
| | | CL | ROM 27484 | |
| | | TL | ROM 27614 | |
| | | LF | ROM 27659 | |
| | | LM | ROM 27734 | |

### 2. Principal component analysis

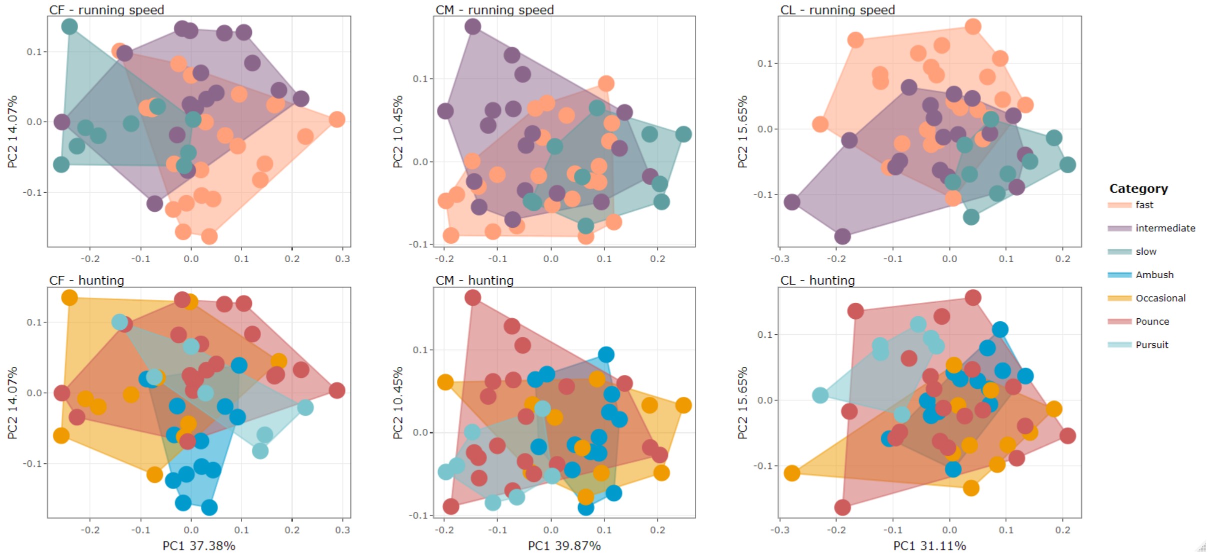

Suppl. Figure 1. Principal component analysis for single cervical vertebrae (CF, CM, CL) with running speed and hunting behaviour morphospaces. For details on the specimens see interactive plot:

https://juliaaschwab.github.io/Vertebrae_locomotion_plots/PCA_cervicals_individual.html

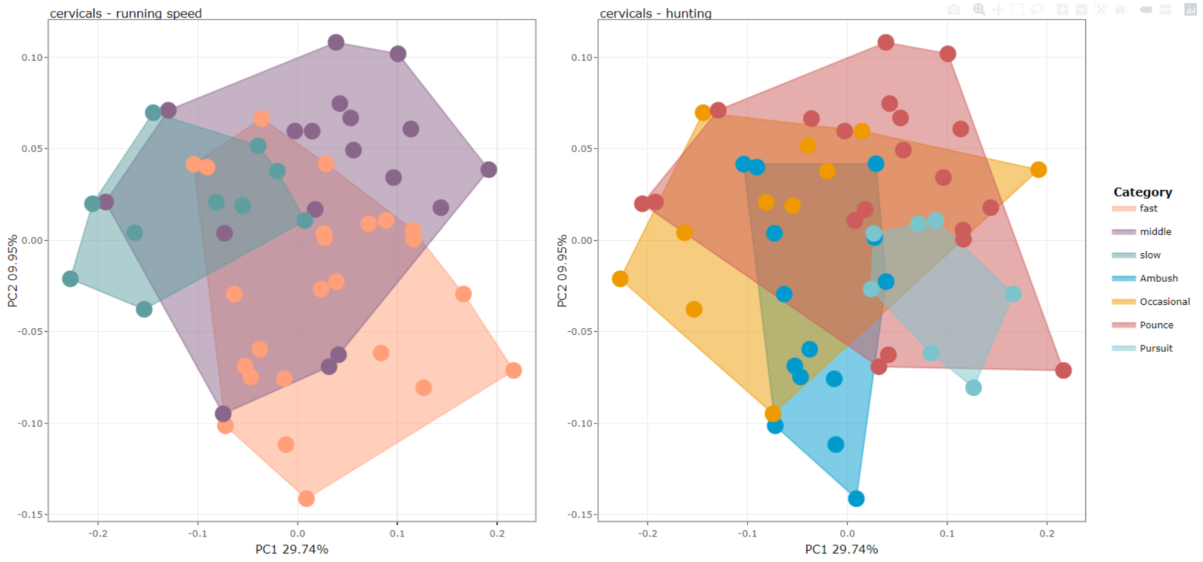

Suppl. Figure 2. Principal component analysis for combined cervical vertebrae with running speed and hunting behaviour morphospaces. For details on the specimens see interactive plot:

https://juliaaschwab.github.io/Vertebrae_locomotion_plots/PCA_cervicals_all.html

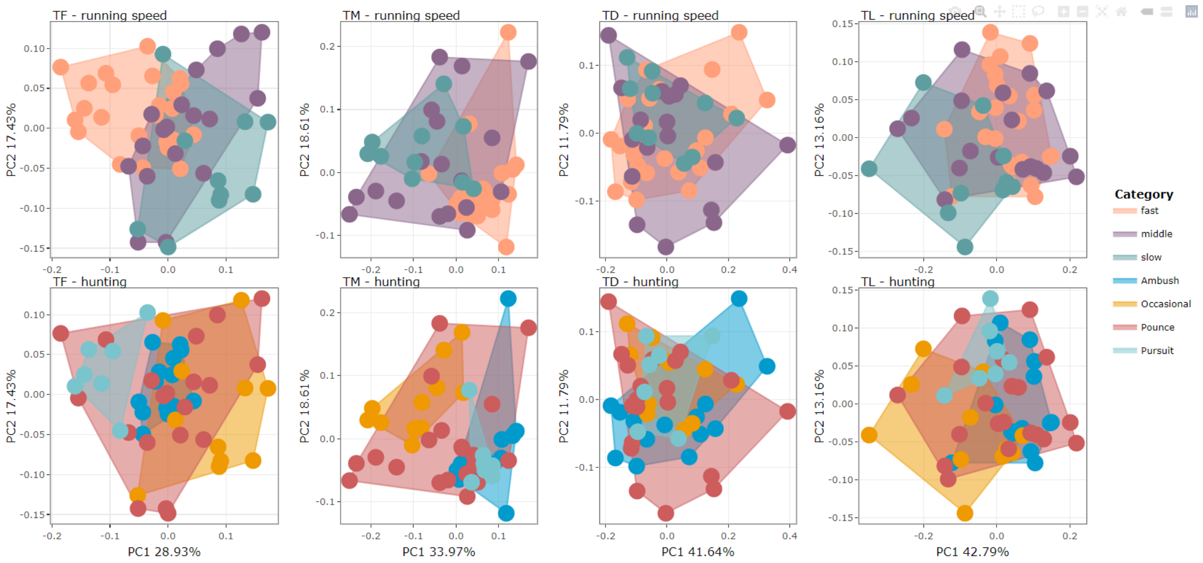

Suppl. Figure 3. Principal component analysis for single thoracic vertebrae (TF, TM, TD, TL) with running speed and hunting behaviour morphospaces. For details on the specimens see interactive plot:

https://juliaaschwab.github.io/Vertebrae_locomotion_plots/PCA_thoracics_individual.html

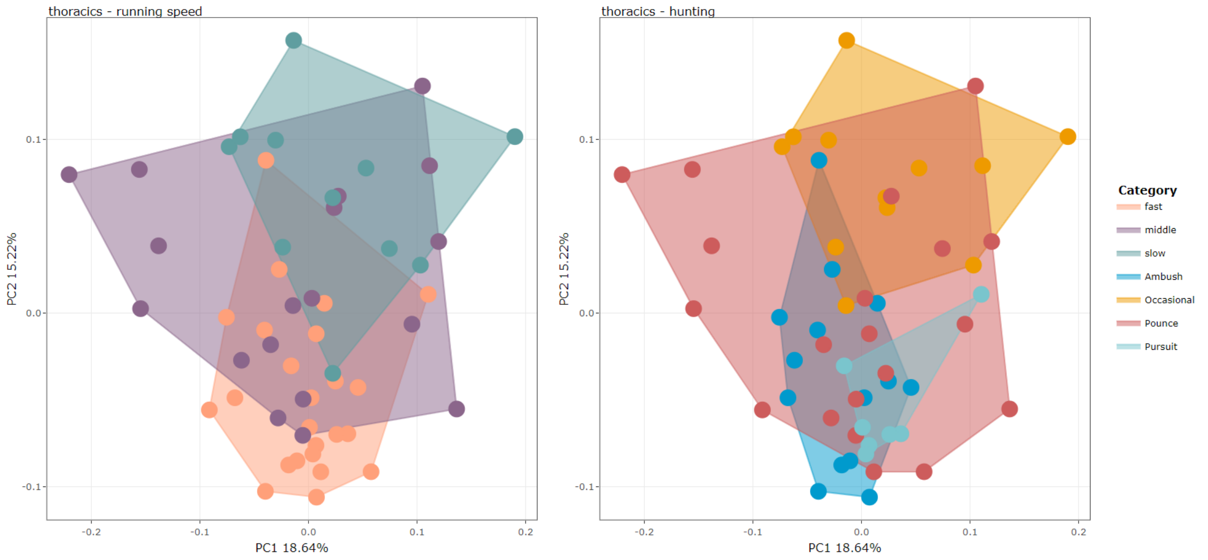

Suppl. Figure 4. Principal component analysis for combined thoracic vertebrae with running speed and hunting behaviour morphospaces. For details on the specimens see interactive plot:

https://juliaaschwab.github.io/Vertebrae_locomotion_plots/PCA_thoracics_all.html

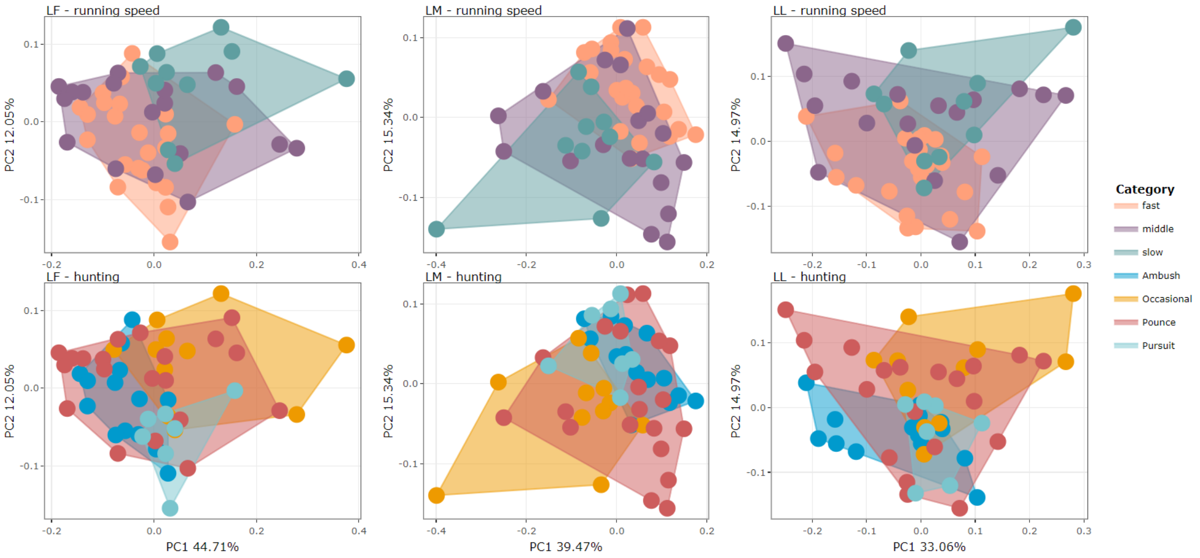

Suppl. Figure 5. Principal component analysis for single lumbar vertebrae (LF, LM, LL) with running speed and hunting behaviour morphospaces. For details on the specimens see interactive plot:

https://juliaaschwab.github.io/Vertebrae_locomotion_plots/PCA_lumbars_individual.html

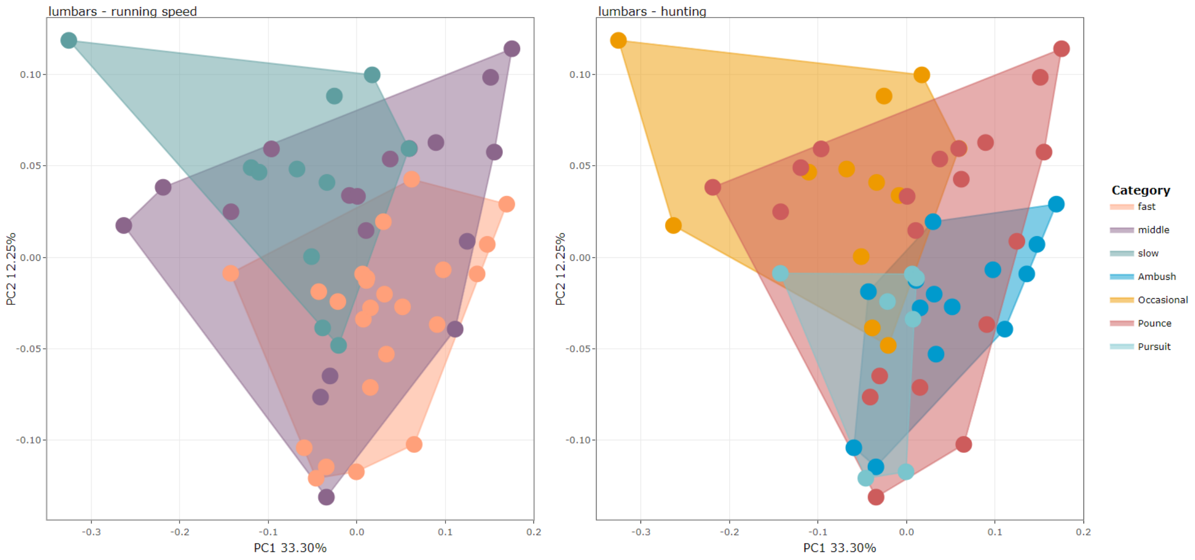

Suppl. Figure 6. Principal component analysis for combined lumbar vertebrae with running speed and hunting behaviour morphospaces. For details on the specimens see interactive plot:

https://juliaaschwab.github.io/Vertebrae_locomotion_plots/PCA_lumbars_all.html

4. **Morphospace clustering analysis**

Suppl. Table 3. Results of the PERMANOVA, for the cervical region, to test which running speeds/hunting behaviours are statistically different. Highlighted in bold are p-value ≤ 0.005.

| **Region** | **Ecology** | **pair** | **R2** | **p value** | **F** |
|---|---|---|---|---|---|
| **cervical** | Hunting | Ambush vs Pounce | 0.139016 | **0.001** | 5.166771 |
| **cervical** | Hunting | Occasional vs Ambush | 0.127488 | 0.011 | 3.360668 |
| **cervical** | Hunting | Occasional vs Pounce | 0.113534 | 0.02 | 3.714179 |
| **cervical** | Hunting | Pursuit vs Ambush | 0.310022 | **0.001** | 8.53709 |
| **cervical** | Hunting | Pursuit vs Occasional | 0.275794 | 0.008 | 6.093147 |
| **cervical** | Hunting | Pursuit vs Pounce | 0.086615 | 0.066 | 2.370725 |
| **cervical** | Running | fast vs intermediate | 0.07824 | 0.009 | 3.395254 |
| **cervical** | Running | fast vs slow | 0.227979 | **0.001** | 9.449665 |
| **cervical** | Running | slow vs intermediate | 0.208826 | 0.007 | 6.862553 |
| **CF** | Hunting | Ambush vs Pounce | 0.125997 | 0.007 | 4.613128 |
| **CF** | Hunting | Occasional vs Ambush | 0.170926 | **0.005** | 4.741785 |
| **CF** | Hunting | Occasional vs Pounce | 0.123496 | 0.022 | 4.086001 |
| **CF** | Hunting | Pursuit vs Ambush | 0.145692 | 0.011 | 3.240211 |
| **CF** | Hunting | Pursuit vs Occasional | 0.179903 | 0.023 | 3.509888 |
| **CF** | Hunting | Pursuit vs Pounce | 0.050552 | 0.241 | 1.331092 |
| **CF** | Running | fast vs intermediate | 0.065112 | 0.034 | 2.785865 |
| **CF** | Running | fast vs slow | 0.225585 | **0.001** | 9.321511 |
| **CF** | Running | slow vs intermediate | 0.219024 | **0.001** | 7.291666 |
| **CM** | Hunting | Ambush vs Pounce | 0.143972 | **0.003** | 5.381971 |
| **CM** | Hunting | Occasional vs Ambush | 0.0351 | 0.488 | 0.836665 |
| **CM** | Hunting | Occasional vs Pounce | 0.105914 | 0.016 | 3.435341 |
| **CM** | Hunting | Pursuit vs Ambush | 0.395244 | **0.001** | 12.41764 |
| **CM** | Hunting | Pursuit vs Occasional | 0.253499 | 0.006 | 5.433338 |
| **CM** | Hunting | Pursuit vs Pounce | 0.063923 | 0.142 | 1.707192 |
| **CM** | Running | fast vs intermediate | 0.048327 | 0.088 | 2.031233 |
| **CM** | Running | fast vs slow | 0.132043 | 0.008 | 4.868188 |
| **CM** | Running | slow vs intermediate | 0.166148 | 0.009 | 5.18059 |
| **CL** | Hunting | Ambush vs Pounce | 0.082345 | 0.026 | 2.871498 |
| **CL** | Hunting | Occasional vs Ambush | 0.131579 | **0.005** | 3.484854 |
| **CL** | Hunting | Occasional vs Pounce | 0.060313 | 0.127 | 1.861339 |
| **CL** | Hunting | Pursuit vs Ambush | 0.263981 | **0.001** | 6.814564 |
| **CL** | Hunting | Pursuit vs Occasional | 0.275476 | **0.002** | 6.083479 |
| **CL** | Hunting | Pursuit vs Pounce | 0.112608 | 0.022 | 3.172443 |
| **CL** | Running | fast vs intermediate | 0.083868 | 0.009 | 3.661816 |
| **CL** | Running | fast vs slow | 0.226501 | **0.001** | 9.370442 |

| | | | | | |
|---|---|---|---|---|---|
| **CL** | Running | slow vs intermediate | 0.15243 | **0.002** | 4.675918 |

Suppl. Table 4. Results of the PERMANOVA, for thoracic vertebrae, to test which running speeds/hunting behaviours are statistically different. Highlighted in bold are p-value ≤ 0.005.

| Region | Ecology | pair | R2 | p value | F |
|---|---|---|---|---|---|
| **thoracic** | Hunting | Ambush vs Pounce | 0.078255 | 0.029 | 2.716768 |
| **thoracic** | Hunting | Occasional vs Ambush | 0.278319 | **0.001** | 9.255673 |
| **thoracic** | Hunting | Occasional vs Pounce | 0.100979 | 0.026 | 3.369615 |
| **thoracic** | Hunting | Pursuit vs Ambush | 0.171996 | 0.014 | 3.946736 |
| **thoracic** | Hunting | Pursuit vs Occasional | 0.368755 | **0.001** | 9.930904 |
| **thoracic** | Hunting | Pursuit vs Pounce | 0.105533 | 0.018 | 2.949614 |
| **thoracic** | Running | fast vs intermediate | 0.088541 | **0.004** | 3.885702 |
| **thoracic** | Running | fast vs slow | 0.244532 | **0.001** | 10.68151 |
| **thoracic** | Running | slow vs intermediate | 0.057123 | 0.167 | 1.635766 |
| **TF** | Hunting | Ambush vs Pounce | 0.060577 | 0.085 | 2.063468 |
| **TF** | Hunting | Occasional vs Ambush | 0.221011 | **0.001** | 6.525452 |
| **TF** | Hunting | Occasional vs Pounce | 0.077747 | 0.068 | 2.444736 |
| **TF** | Hunting | Pursuit vs Ambush | 0.328981 | **0.001** | 9.315164 |
| **TF** | Hunting | Pursuit vs Occasional | 0.436698 | **0.001** | 12.40395 |
| **TF** | Hunting | Pursuit vs Pounce | 0.154511 | **0.002** | 4.568688 |
| **TF** | Running | fast vs intermediate | 0.127879 | **0.001** | 5.865196 |
| **TF** | Running | fast vs slow | 0.257677 | **0.001** | 11.10791 |
| **TF** | Running | slow vs intermediate | 0.059245 | 0.188 | 1.637369 |
| **TM** | Hunting | Ambush vs Pounce | 0.1186 | **0.002** | 4.305883 |
| **TM** | Hunting | Occasional vs Ambush | 0.354301 | **0.001** | 12.62032 |
| **TM** | Hunting | Occasional vs Pounce | 0.114479 | 0.019 | 3.749096 |
| **TM** | Hunting | Pursuit vs Ambush | 0.046211 | 0.483 | 0.920544 |
| **TM** | Hunting | Pursuit vs Occasional | 0.414542 | **0.001** | 11.32901 |
| **TM** | Hunting | Pursuit vs Pounce | 0.116093 | 0.032 | 3.283521 |
| **TM** | Running | fast vs intermediate | 0.143776 | **0.001** | 6.716743 |
| **TM** | Running | fast vs slow | 0.283849 | **0.001** | 12.68329 |
| **TM** | Running | slow vs intermediate | 0.023128 | 0.63 | 0.615569 |
| **TD** | Hunting | Ambush vs Pounce | 0.022204 | 0.527 | 0.726648 |
| **TD** | Hunting | Occasional vs Ambush | 0.065129 | 0.171 | 1.602322 |
| **TD** | Hunting | Occasional vs Pounce | 0.045349 | 0.239 | 1.377578 |
| **TD** | Hunting | Pursuit vs Ambush | 0.075913 | 0.186 | 1.560827 |
| **TD** | Hunting | Pursuit vs Occasional | 0.164102 | 0.027 | 3.1411 |
| **TD** | Hunting | Pursuit vs Pounce | 0.032277 | 0.477 | 0.833834 |
| **TD** | Running | fast vs intermediate | 0.016538 | 0.559 | 0.672662 |
| **TD** | Running | fast vs slow | 0.058016 | 0.143 | 1.970849 |
| **TD** | Running | slow vs intermediate | 0.030044 | 0.488 | 0.805329 |
| **TL** | Hunting | Ambush vs Pounce | 0.05048 | 0.147 | 1.701221 |

| | | | | | |
|---|---|---|---|---|---|
| **TL** | Hunting | Occasional vs Ambush | 0.226724 | **0.005** | 6.743596 |
| **TL** | Hunting | Occasional vs Pounce | 0.100982 | 0.042 | 3.257409 |
| **TL** | Hunting | Pursuit vs Ambush | 0.193976 | **0.003** | 4.572488 |
| **TL** | Hunting | Pursuit vs Occasional | 0.176836 | 0.017 | 3.437201 |
| **TL** | Hunting | Pursuit vs Pounce | 0.081578 | 0.1 | 2.220606 |
| **TL** | Running | fast vs intermediate | 0.028804 | 0.29 | 1.186322 |
| **TL** | Running | fast vs slow | 0.191481 | **0.002** | 7.578515 |
| **TL** | Running | slow vs intermediate | 0.091986 | 0.09 | 2.633932 |

Suppl. Table 5. Results of the PERMANOVA, for lumbar vertebrae, to test which running speeds/hunting behaviours are statistically different. Highlighted in bold are p-value ≤ 0.005.

| **Region** | **Ecology** | **pair** | **R2** | **p value** | **F** |
|---|---|---|---|---|---|
| **lumbar** | Hunting | Ambush vs Pounce | 0.047002 | 0.167 | 1.578237 |
| **lumbar** | Hunting | Occasional vs Ambush | 0.278492 | **0.001** | 9.263671 |
| **lumbar** | Hunting | Occasional vs Pounce | 0.090731 | 0.031 | 2.993546 |
| **lumbar** | Hunting | Pursuit vs Ambush | 0.132869 | 0.051 | 2.911332 |
| **lumbar** | Hunting | Pursuit vs Occasional | 0.159801 | 0.015 | 3.233303 |
| **lumbar** | Hunting | Pursuit vs Pounce | 0.067226 | 0.123 | 1.801773 |
| **lumbar** | Running | fast vs intermediate | 0.07065 | 0.029 | 3.040822 |
| **lumbar** | Running | fast vs slow | 0.231331 | **0.001** | 9.931331 |
| **lumbar** | Running | slow vs intermediate | 0.056503 | 0.16 | 1.616941 |
| **LF** | Hunting | Ambush vs Pounce | 0.04803 | 0.18 | 1.614505 |
| **LF** | Hunting | Occasional vs Ambush | 0.261381 | **0.002** | 8.139177 |
| **LF** | Hunting | Occasional vs Pounce | 0.100427 | 0.051 | 3.237515 |
| **LF** | Hunting | Pursuit vs Ambush | 0.232573 | **0.001** | 5.758065 |
| **LF** | Hunting | Pursuit vs Occasional | 0.169818 | 0.017 | 3.272871 |
| **LF** | Hunting | Pursuit vs Pounce | 0.101278 | 0.055 | 2.817278 |
| **LF** | Running | fast vs intermediate | 0.034694 | 0.215 | 1.437644 |
| **LF** | Running | fast vs slow | 0.230776 | **0.001** | 9.600363 |
| **LF** | Running | slow vs intermediate | 0.091635 | 0.076 | 2.622871 |
| **LM** | Hunting | Ambush vs Pounce | 0.057256 | 0.1 | 1.943479 |
| **LM** | Hunting | Occasional vs Ambush | 0.243805 | **0.001** | 7.415428 |
| **LM** | Hunting | Occasional vs Pounce | 0.125782 | 0.016 | 4.172502 |
| **LM** | Hunting | Pursuit vs Ambush | 0.092067 | 0.111 | 1.926652 |
| **LM** | Hunting | Pursuit vs Occasional | 0.141857 | 0.045 | 2.644903 |
| **LM** | Hunting | Pursuit vs Pounce | 0.092546 | 0.059 | 2.549598 |
| **LM** | Running | fast vs intermediate | 0.076321 | 0.037 | 3.305099 |
| **LM** | Running | fast vs slow | 0.197623 | **0.001** | 7.881526 |
| **LM** | Running | slow vs intermediate | 0.06884 | 0.14 | 1.922156 |
| **LL** | Hunting | Ambush vs Pounce | 0.052643 | 0.122 | 1.778202 |
| **LL** | Hunting | Occasional vs Ambush | 0.206071 | **0.002** | 5.969839 |
| **LL** | Hunting | Occasional vs Pounce | 0.039429 | 0.291 | 1.190386 |

| | | | | | |
|---|---|---|---|---|---|
| **LL** | Hunting | Pursuit vs Ambush | 0.074327 | 0.198 | 1.525608 |
| **LL** | Hunting | Pursuit vs Occasional | 0.112988 | 0.109 | 2.038097 |
| **LL** | Hunting | Pursuit vs Pounce | 0.029111 | 0.587 | 0.749594 |
| **LL** | Running | fast vs intermediate | 0.072948 | 0.026 | 3.147505 |
| **LL** | Running | fast vs slow | 0.140687 | **0.001** | 5.239065 |
| **LL** | Running | slow vs intermediate | 0.017023 | 0.76 | 0.450276 |

Suppl. Table 6. Results of the PERMANOVA, for the full column, to test which running speeds/hunting behaviours are statistically different. Highlighted in bold are p-value ≤ 0.005.

| **Region** | **Ecology** | **pair** | **R2** | **p value** | **F** |
|---|---|---|---|---|---|
| **all** | Hunting | Pursuit vs Occasional | 0.303078 | **0.001** | 6.958079 |
| **all** | Hunting | Pursuit vs Ambush | 0.246049 | **0.001** | 6.200567 |
| **all** | Hunting | Occasional vs Ambush | 0.271727 | **0.001** | 8.581563 |
| **all** | Running | fast vs slow | 0.26945 | **0.001** | 11.80261 |
| **all** | Running | fast vs intermediate | 0.095217 | **0.003** | 4.209475 |
| **all** | Hunting | Ambush vs Pounce | 0.10963 | **0.004** | 3.940113 |
| **all** | Hunting | Occasional vs Pounce | 0.115309 | 0.013 | 3.779805 |
| **all** | Running | slow vs intermediate | 0.116364 | 0.02 | 3.423889 |
| **all** | Hunting | Pursuit vs Pounce | 0.09192 | 0.048 | 2.530618 |

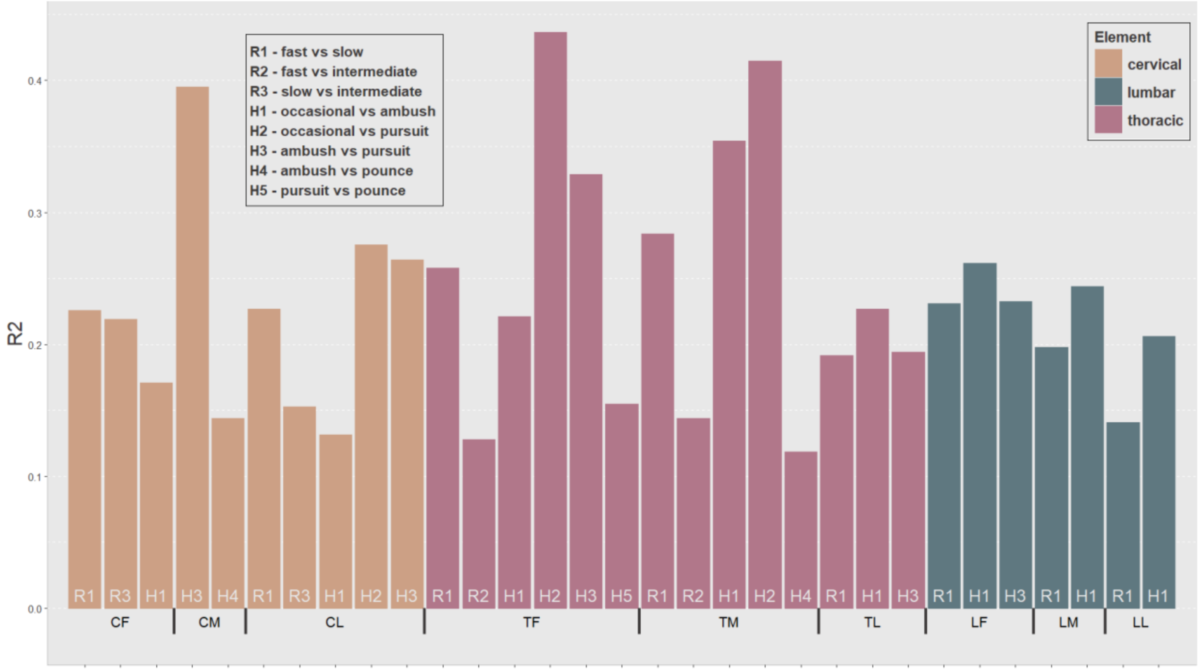

Suppl. Figure 7. PERMANOVA for individual vertebrae. Only significant (p-value ≤ 0.005) results are shown.

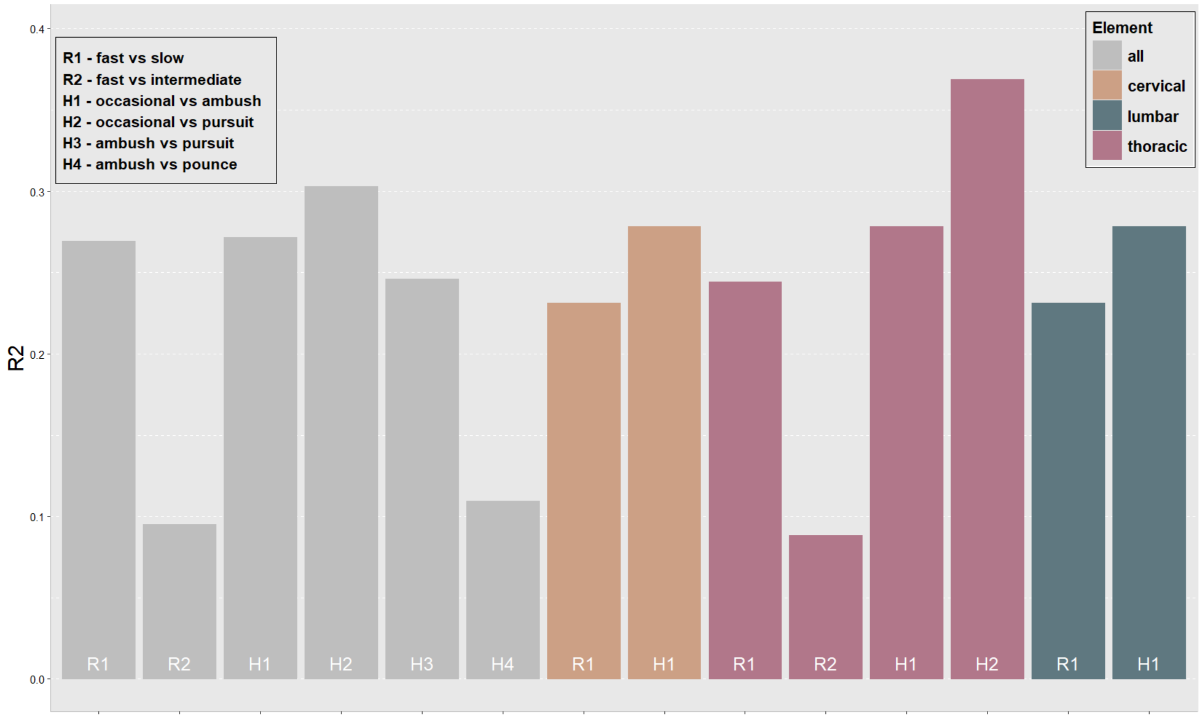

Suppl. Figure 8. PERMANOVA for combined vertebrae. Only significant (p-value ≤ 0.005) results are shown.

5. **pGLS regressions**

Suppl. Table 7. Results of the PGLS, for cervical vertebrae, to test for correlation which running speeds, hunting behaviours and centroid size. Highlighted in bold are p-value ≤ 0.005 and the highest $R^2$.

| Region | pair | $R^2$ | p-value |
|---|---|---|---|
| **CF** | Csize | 0.07643 | 0.02 |
| **CF** | hunting | 0.13738 | 0.024 |
| **CF** | running | 0.17483 | **0.001** |
| **CM** | Csize | 0.07907 | 0.018 |
| **CM** | hunting | 0.1261 | 0.053 |
| **CM** | running | 0.17224 | **0.001** |
| **CL** | Csize | 0.08256 | 0.015 |
| **CL** | hunting | 0.12341 | 0.069 |
| **CL** | running | 0.16167 | **0.001** |
| **cervical** | hunting | 0.11226 | 0.088 |
| **cervical** | running | 0.13475 | **0.004** |

Suppl. Table 8. Results of the PGLS, for thoracic vertebrae, to test for correlation which running speeds, hunting behaviours and centroid size. Highlighted in bold are p-value ≤ 0.005 and the highest $R^2$.

| Region | pair | $R^2$ | p-value |
|---|---|---|---|
| **TF** | Csize | 0.07688 | 0.027 |
| **TF** | hunting | 0.14391 | 0.028 |
| **TF** | running | 0.17998 | **0.001** |
| **TM** | Csize | 0.08063 | 0.019 |
| **TM** | hunting | 0.13601 | 0.039 |
| **TM** | running | 0.15574 | 0.006 |
| **TD** | Csize | 0.06481 | 0.044 |
| **TD** | hunting | 0.12573 | 0.056 |
| **TD** | running | 0.1348 | 0.008 |
| **TL** | Csize | 0.10461 | 0.009 |
| **TL** | hunting | 0.1527 | 0.016 |
| **TL** | running | 0.1718 | **0.002** |
| **thoracic** | hunting | 0.13908 | 0.021 |
| **thoracic** | running | 0.16697 | **0.001** |

Suppl. Table 9. Results of the PGLS, for lumbar vertebrae, to test for correlation which running speeds, hunting behaviours and centroid size. Highlighted in bold are p-value ≤ 0.005 and the highest $R^2$.

| Region | pair | $R^2$ | p-value |
|---|---|---|---|
| **LF** | Csize | 0.10996 | 0.009 |
| **LF** | hunting | 0.17163 | 0.005 |
| **LF** | running | 0.18183 | **0.001** |
| **LM** | Csize | 0.10998 | 0.011 |
| **LM** | hunting | 0.13795 | 0.036 |
| **LM** | running | 0.18004 | **0.001** |
| **LL** | Csize | 0.08753 | 0.013 |
| **LL** | hunting | 0.13358 | 0.024 |
| **LL** | running | 0.14934 | **0.002** |
| **lumbar** | hunting | 0.12842 | 0.041 |
| **lumbar** | running | 0.1611 | **0.001** |

Suppl. Table 10. Results of the PGLS, for all combined vertebrae as well as different tested combinations, to test for correlation which running speeds, hunting behaviours and centroid size. Highlighted in bold are p-value ≤ 0.005 and the highest $R^2$.

| Region | pair | $R^2$ | p-value |
|---|---|---|---|
| **all** | hunting | 0.11226 | 0.088 |
| **all** | running | 0.13475 | **0.004** |
| **CF, CL** | running | 0.13851 | **0.004** |
| **CF, CM** | running | 0.1352 | **0.005** |
| **CF, LF** | running | 0.14957 | **0.001** |
| **CF, LL** | running | 0.14834 | **0.001** |
| **CF, LM** | running | 0.1571 | **0.001** |
| **CF, TD** | running | 0.14016 | **0.001** |
| **CF, TF** | running | 0.15458 | **0.001** |
| **CF, TL** | running | 0.14552 | **0.001** |
| **CF, TM** | running | 0.16178 | **0.001** |
| **CM, CL** | running | 0.13027 | **0.005** |
| **CM, LF** | running | 0.14191 | **0.004** |
| **CM, LL** | running | 0.14144 | **0.001** |
| **CM, LM** | running | 0.15 | **0.001** |
| **CM, TD** | running | 0.13231 | **0.003** |
| **CM, TF** | running | 0.14689 | **0.002** |
| **CM, TL** | running | 0.13785 | **0.004** |
| **CM, TM** | running | 0.15504 | **0.001** |
| **CL, LF** | running | 0.14555 | **0.003** |
| **CL, LL** | running | 0.14468 | **0.001** |
| **CL, LM** | running | 0.15366 | **0.001** |

| | | | |
|---|---|---|---|
| **CL, TD** | running | 0.13571 | **0.002** |
| **CL, TF** | running | 0.15074 | **0.002** |
| **CL, TL** | running | 0.14135 | **0.003** |
| **CL, TM** | running | 0.15869 | **0.001** |
| **TF, LF** | running | 0.16346 | **0.001** |
| **TF, LL** | running | 0.16049 | **0.001** |
| **TF, LM** | running | 0.17182 | **0.001** |
| **TF, TD** | running | 0.15214 | **0.001** |
| **TF, TL** | running | 0.15843 | **0.001** |
| **TF, TM** | running | 0.17693 | **0.001** |
| **TM, LF** | running | 0.17065 | **0.001** |
| **TM, LL** | running | 0.16708 | **0.001** |
| **TM, LM** | running | 0.17837 | **0.001** |
| **TM, TD** | running | 0.15972 | **0.001** |
| **TM, TL** | running | 0.16573 | **0.001** |
| **TD, LF** | running | 0.14708 | **0.001** |
| **TD, LL** | running | 0.14608 | **0.001** |
| **TD, LM** | running | 0.15488 | **0.001** |
| **TD, TL** | running | 0.14298 | **0.002** |
| **TL, LF** | running | 0.153 | **0.002** |
| **TL, LL** | running | 0.15135 | **0.002** |
| **TL, LM** | running | 0.16087 | **0.002** |
| **LF, LL** | running | 0.15542 | **0.002** |
| **LF, LM** | running | 0.1657 | **0.002** |
| **LM, LL** | running | 0.16263 | **0.001** |
| **TF, LF, LM** | running | 0.16699 | **0.001** |
| **TF, TL, LF** | running | 0.15816 | **0.002** |
| **TF, TL, LM** | running | 0.1636 | **0.001** |
| **TF, TM, LF** | running | 0.17041 | **0.001** |
| **TF, TM, LM** | running | 0.17577 | **0.001** |
| **TM, LF, LM** | running | 0.17162 | **0.001** |
| **TL, LF, LM** | running | 0.15982 | **0.002** |
| **TF, TL, LF, LM** | running | 0.16211 | **0.002** |
| **TF, TM, LF, LM** | running | 0.17122 | **0.001** |

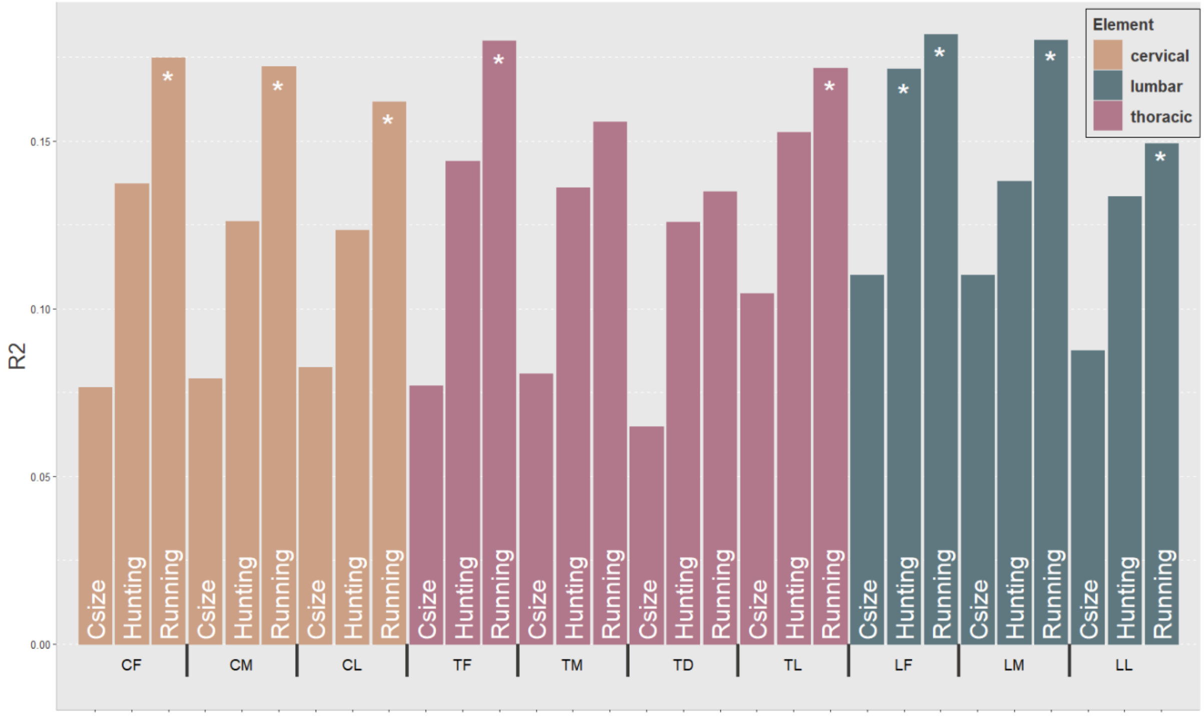

Suppl. Figure 9. PGLS for single vertebrae. Significant (p-value ≤ 0.005) results are shown with an asterisk (*).

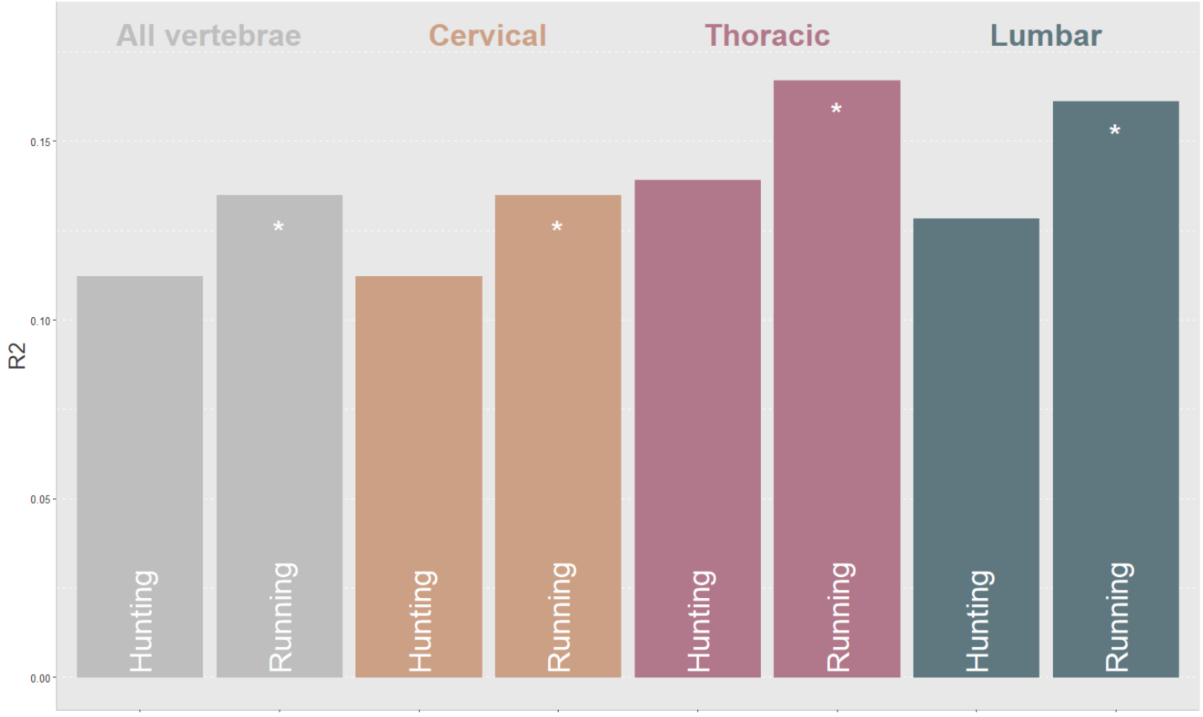

Suppl. Figure 10. PGLS for combined vertebrae. Significant (p-value ≤ 0.005) results are shown with an asterisk (*).

### 6. Canonical variate analysis

Suppl. Table 11. Overall classification accuracy of different vertebrae for running speed based on a canonical variate analysis (CVA).

| CF | fast | intermediate | slow | overall classification accuracy: 76.74419 % |
|---|---|---|---|---|
| **fast** | 89.4737 | 5.2632 | 5.2632 | |
| **intermediate** | 33.3333 | 66.6667 | 0 | |
| **slow** | 0 | 33.3333 | 66.6667 | |
| **CM** | **fast** | **intermediate** | **slow** | overall classification accuracy: 53.48837 % |
| **fast** | 57.8947 | 36.8421 | 5.2632 | |
| **intermediate** | 26.6667 | 46.6667 | 26.6667 | |
| **slow** | 11.1111 | 33.3333 | 55.5556 | |
| **CL** | **fast** | **intermediate** | **slow** | overall classification accuracy: 53.48837 % |
| **fast** | 78.95 | 15.79 | 5.26 | |
| **intermediate** | 33.33 | 26.67 | 40.00 | |
| **slow** | 11.11 | 44.44 | 44.44 | |
| **TL** | **fast** | **intermediate** | **slow** | overall classification accuracy: 55.81395 % |
| **fast** | 73.684 | 10.526 | 15.789 | |
| **intermediate** | 13.333 | 53.333 | 33.333 | |
| **slow** | 22.222 | 55.556 | 22.222 | |
| **LF** | **fast** | **intermediate** | **slow** | overall classification accuracy: 60.46512 % |
| **fast** | 68.4211 | 26.3158 | 5.2632 | |
| **intermediate** | 26.6667 | 53.3333 | 20 | |
| **slow** | 11.1111 | 33.3333 | 55.5556 | |
| **LM** | **fast** | **intermediate** | **slow** | |
| **fast** | 68.421 | 31.579 | 0 | overall classification accuracy: 51.16279 % |
| **intermediate** | 53.333 | 26.667 | 20 | |
| **slow** | 11.111 | 33.333 | 55.556 | |

Suppl. Table 12. Overall classification accuracy of different vertebrae for hunting behaviour based on a canonical variate analysis (CVA).

| LF | Ambush | Occasional | Pounce | Pursuit | |
|---|---|---|---|---|---|
| **Ambush** | 58.33 | 8.33 | 16.67 | 16.67 | overall classification accuracy: 37.2093 % |
| **Occasional** | 10.00 | 50.00 | 30.00 | 10.00 | |
| **Pounce** | 25.00 | 43.75 | 18.75 | 12.50 | |
| **Pursuit** | 40.00 | 0.00 | 40.00 | 20.00 | |

Suppl. Table 13. Overall classification accuracy of all preserved vertebrae (CF, CM, CL, TL, LF and LM) of *Canis dirus* for running speed based on a canonical variate analysis (CVA).

| **All** | **fast** | **intermediate** | **slow** | |
|---|---|---|---|---|
| **fast** | 52.6316 | 42.1053 | 5.2632 | overall classification accuracy: 34.88372 % |
| **intermediate** | 60.0000 | 26.6667 | 13.3333 | |
| **slow** | 33.3333 | 55.5556 | 11.1111 | |

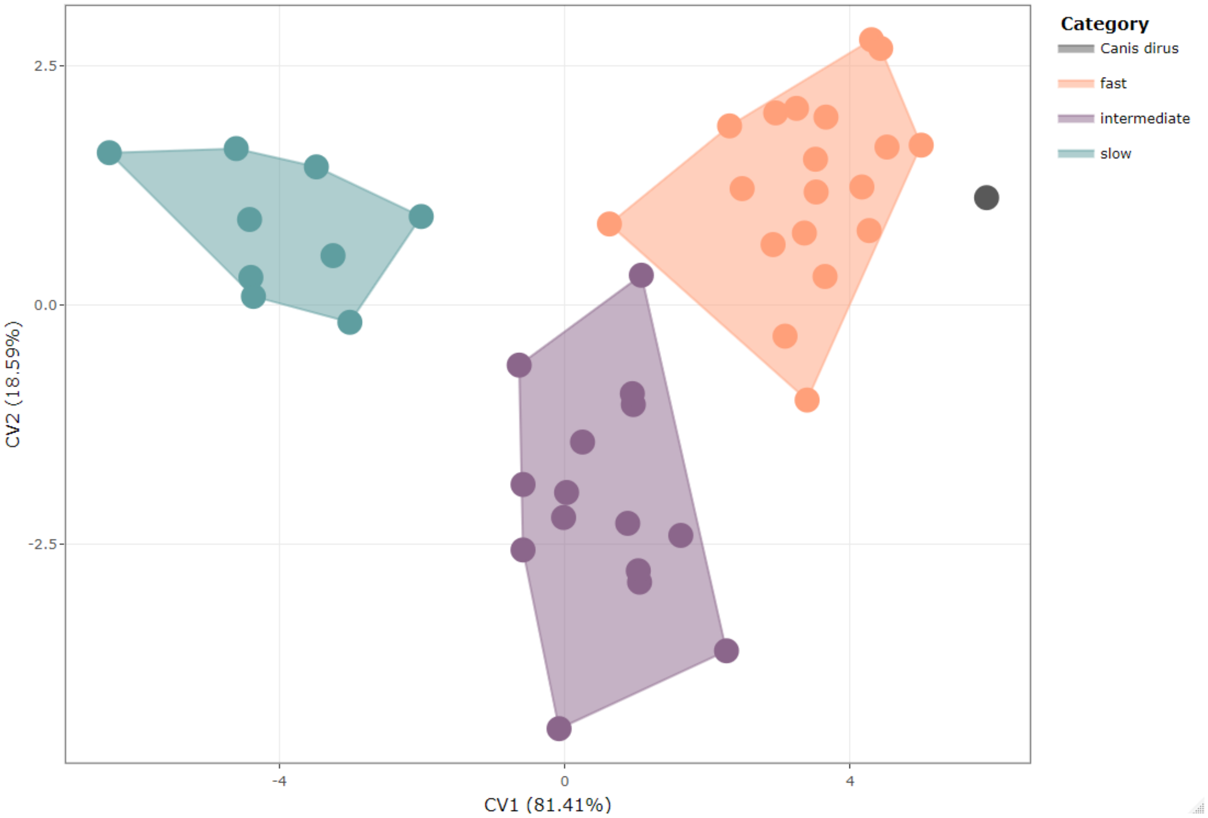

Suppl. Figure 11. CVA of CF and running speed. For details on the specimens see interactive plot:

https://juliaaschwab.github.io/Vertebrae_locomotion_plots/CVA_CF.html

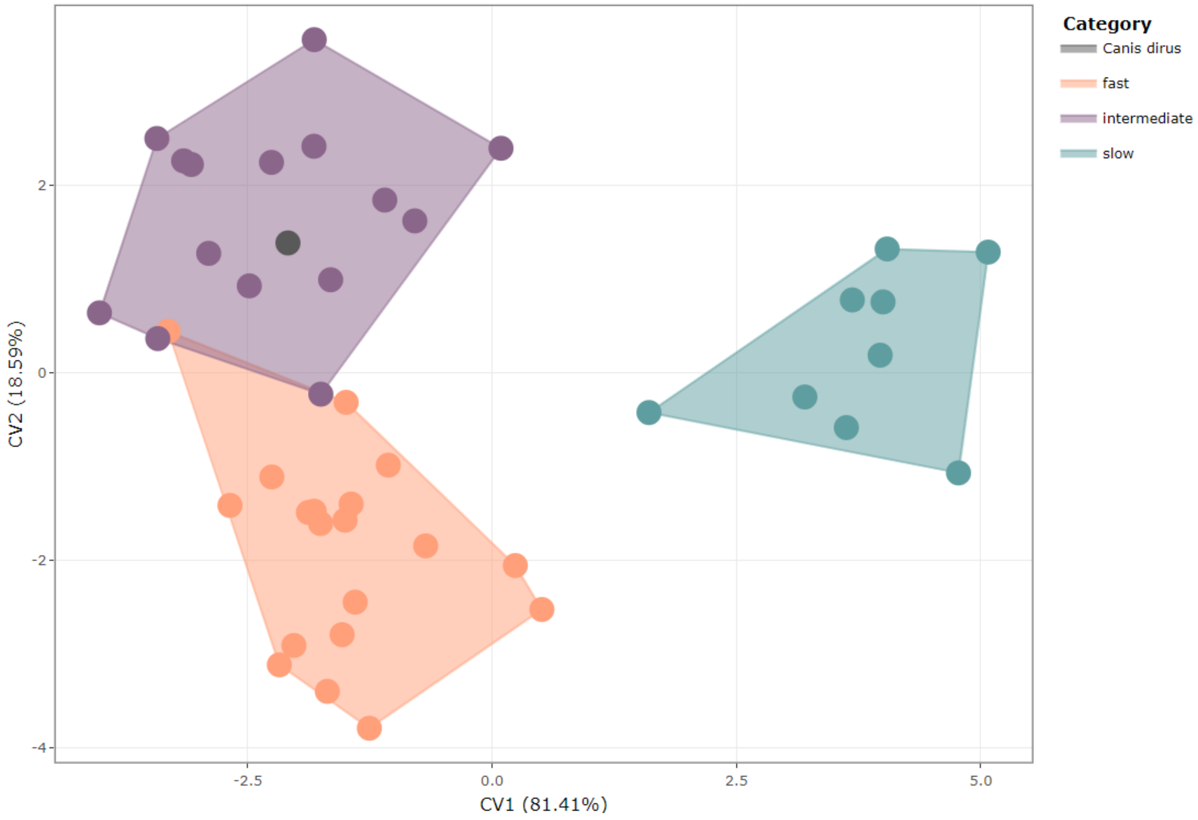

Suppl. Figure 12. CVA of CM and running speed. For details on the specimens see interactive plot:

https://juliaaschwab.github.io/Vertebrae_locomotion_plots/CVA_CM.html

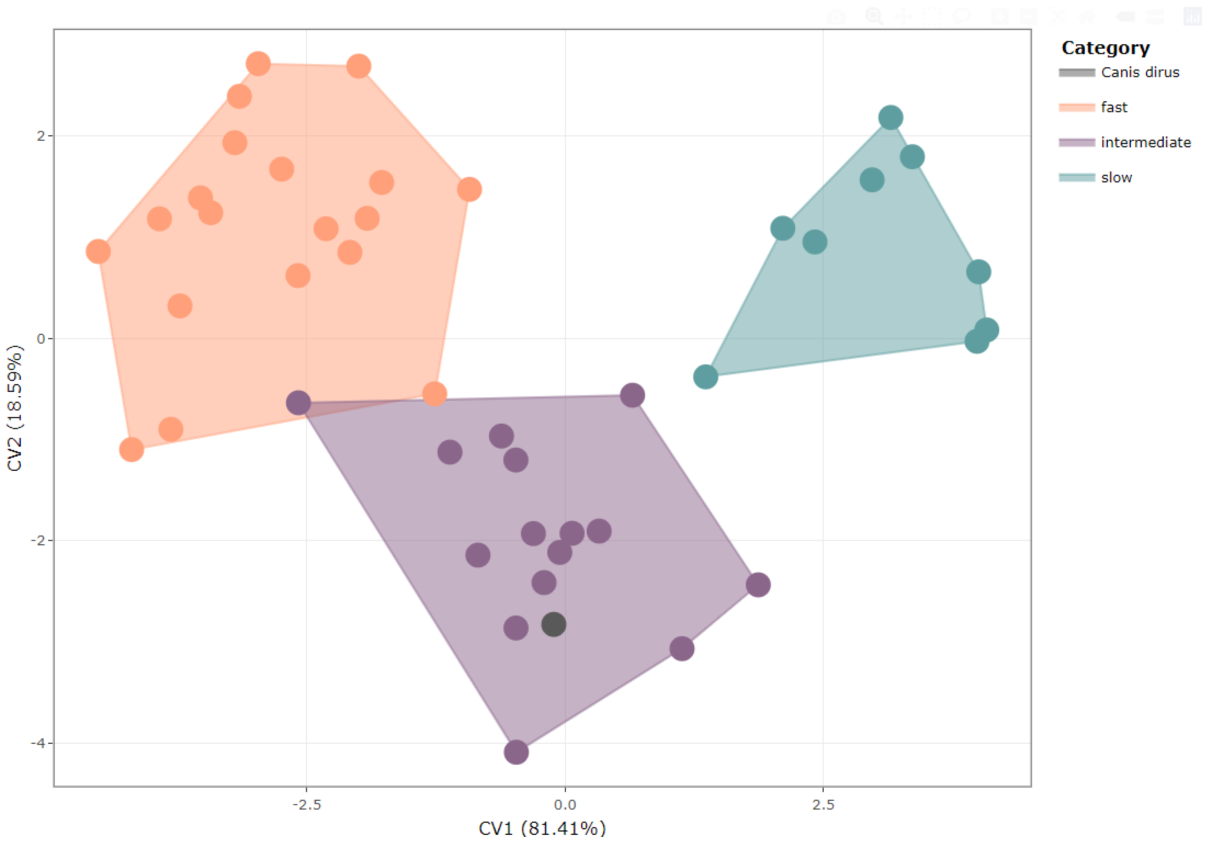

Suppl. Figure 13. CVA of CL and running speed. For details on the specimens see interactive plot:

https://juliaaschwab.github.io/Vertebrae_locomotion_plots/CVA_CL.html

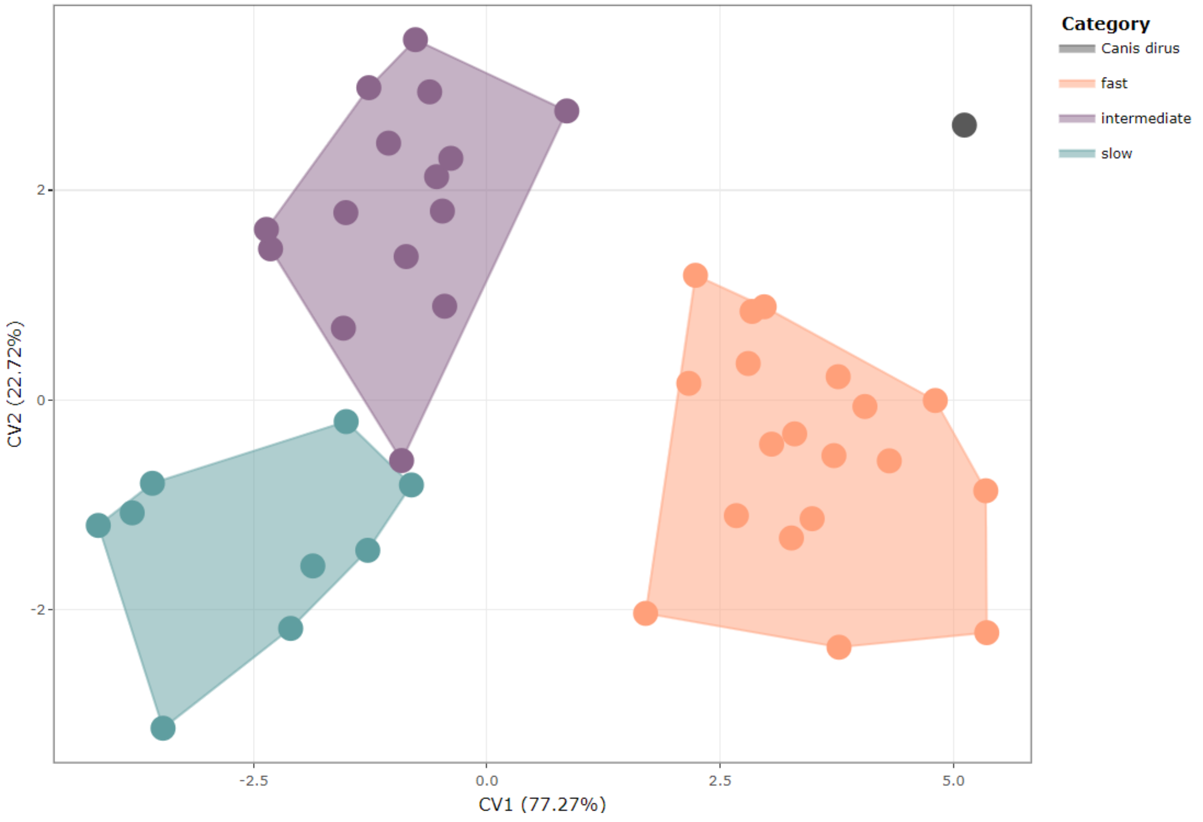

Suppl. Figure 14. CVA of TL and running speed. For details on the specimens see interactive plot:

https://juliaaschwab.github.io/Vertebrae_locomotion_plots/CVA_TL.html

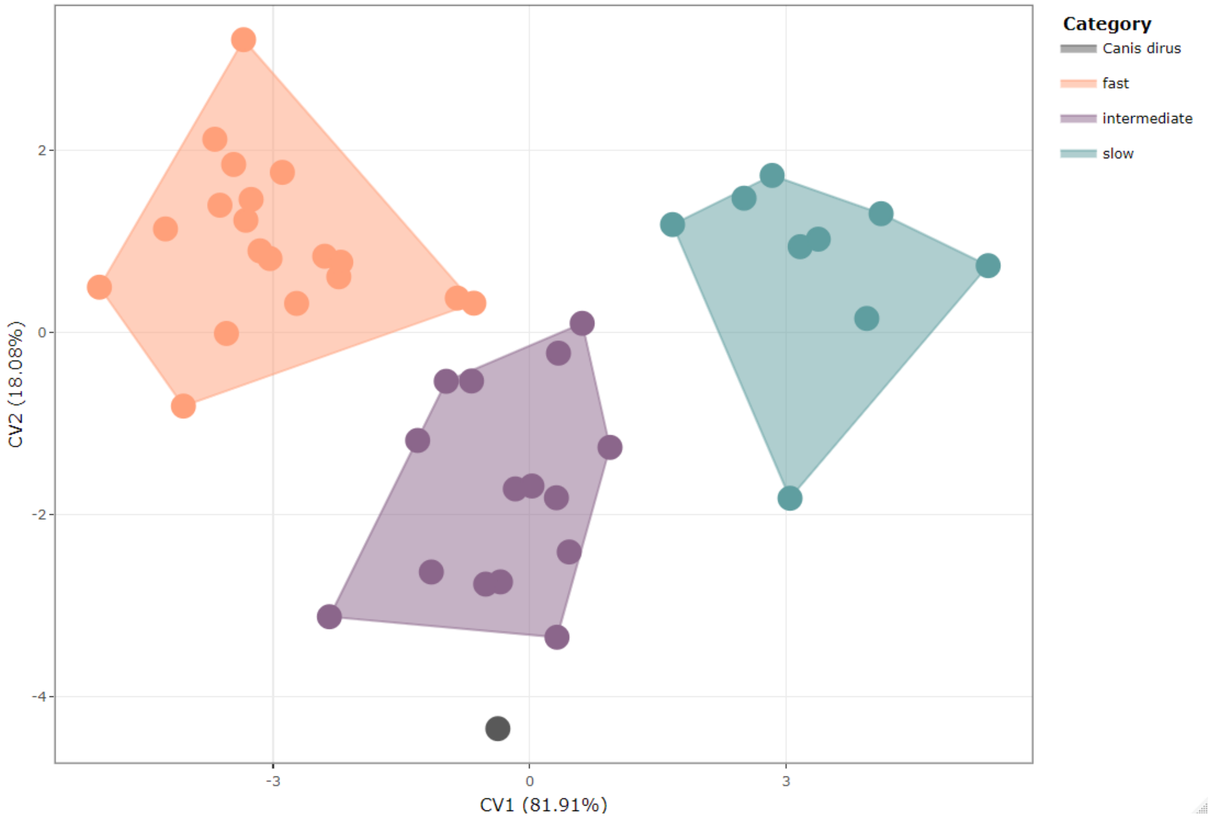

Suppl. Figure 15. CVA of LM and running speed. For details on the specimens see interactive plot:

https://juliaaschwab.github.io/Vertebrae_locomotion_plots/CVA_LM.html

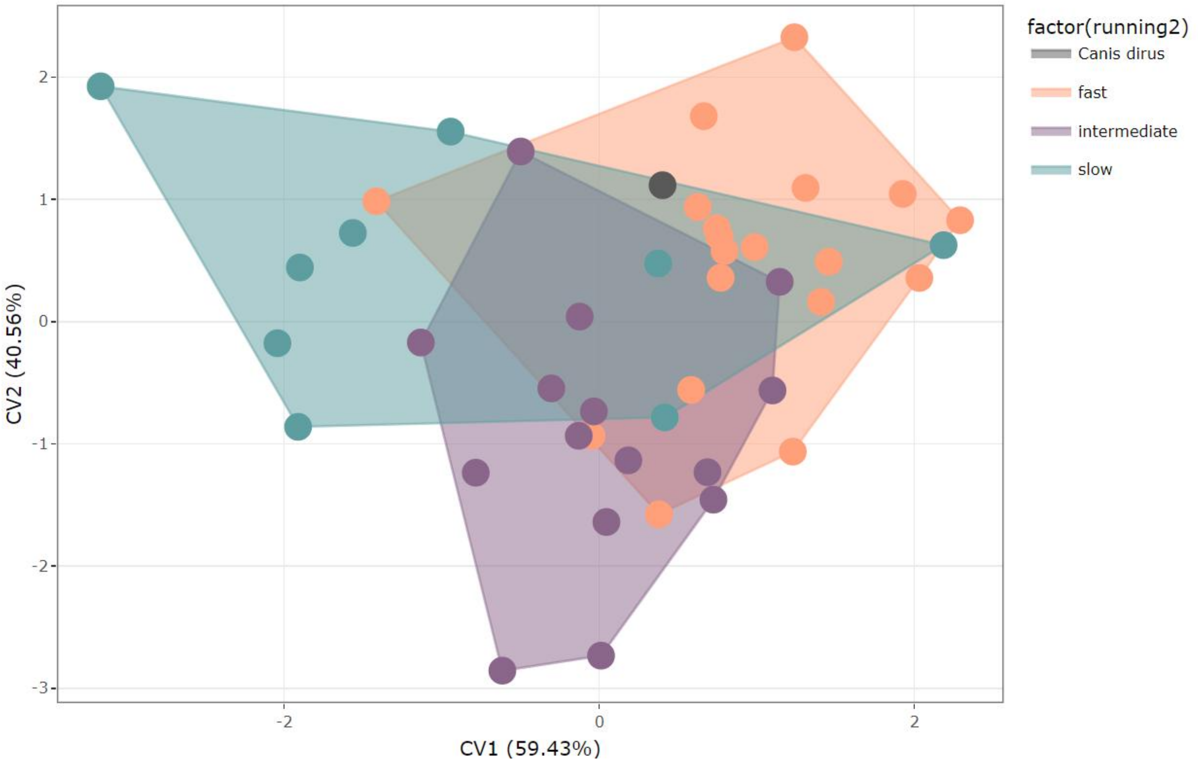

Suppl. Figure 16. CVA of all preserved vertebrae (CF, CM, CL, TL, LF and LM) and running speed. For details on the specimens see interactive plot:

https://juliaaschwab.github.io/Vertebrae_locomotion_plots/CVA_all.html

### 7. Mahalanobis Distances

Suppl. Table 14. Probability of Mahalanobis Distances for hunting behaviour for *Canis dirus*.

| | Ambush | Occasional | Pounce | Pursuit |
|---|---|---|---|---|
| LF | 0.0000 | 0.0000 | 0.0000 | 1.0000 |

Suppl. Table 15. Probability of Mahalanobis Distances for running speed for *Canis dirus*.

| | fast | intermediate | slow |
|---|---|---|---|
| CF | 1.00000 | 0.00000 | 0.00000 |
| CM | 0.00711 | 0.99289 | 0.00000 |
| CL | 0.00012 | 0.99988 | 0.00000 |
| TL | 1.00000 | 0.00000 | 0.00000 |
| LF | 0.99903 | 0.00000 | 0.00096 |
| LM | 0.42353 | 0.57399 | 0.00248 |
| all | 0.79515 | 0.09167 | 0.11318 |
